# Modeling Dynamical Vision with Biologically Plausible Recurrent Convolutional Networks

**DOI:** 10.1101/2025.08.11.669756

**Authors:** Robin Gutzen, Grace W Lindsay

## Abstract

Convolutional Neural Networks (CNNs) trained for image recognition have demonstrated remarkable conceptual similarities to the primate ventral visual pathway, but their standard feedforward architectures lack the recurrent connections that are ubiquitous in visual cortex. Such recurrence is thought to underlie spatiotemporal phenomena including adaptation, delayed normalization, and robustness to noisy input. However, incorporating functionally beneficial recurrence into CNNs that captures spatiotemporal phenomena of biological vision remains challenging. Although recent advances have incorporated neurobiological constraints, the field lacks accessible tools for systematically comparing how different architectural choices, such as recurrence type, temporal delays, and connectivity patterns, shape neural dynamics and behavior. Here, we introduce DynVision, a modular open-source toolbox for constructing and evaluating biologically plausible recurrent convolutional neural networks (RCNNs). DynVision implements numerical ODE solvers with heterogeneous delays, supports five types of lateral recurrence ranging from simple self-connections to cortically-organized local recurrence, and separates scientific modeling decisions from implementation details through a configuration-driven design. Training is computationally efficient, achieving a 52% speedup over reference implementations. We demonstrate the framework through systematic exploration of the parameter space, revealing that qualitative differences in temporal dynamics are highly sensitive to often-implicit modeling choices such as the target location of recurrent integration and the temporal window used for loss computation. Critically, we find that continuous-time recurrent dynamics can naturally give rise to cortical temporal phenomena without requiring explicit divisive normalization, while a different recurrent configuration produces noise robustness approaching human-level performance. These findings suggest functionally distinct configurations of recurrence and highlight the challenge of creating fully realistic models, thus emphasizing the need for a comprehensive and cohesive modeling framework to aid exploration. Code and documentation are available at https://github.com/Lindsay-Lab/DynVision/.

## 1 Introduction

The primate visual system is characterized by abundant recurrent connections [1]. In the ventral stream, lateral connections link neurons within a cortical region and feedback connections project from higher areas such as V4 back to lower ones like V1. These connections are believed to give rise to characteristic temporal response properties observed in cortical recordings and play a crucial role in the system’s remarkable ability to integrate information over time and recognize objects under diverse viewing conditions [1, 2, 3, 4]. Understanding how recurrence shapes these dynamics is central to both computational neuroscience and the development of more brain-like artificial vision systems.

For these reasons, many researchers have attempted to integrate these connections into convolutional neural networks (CNNs), one of the leading classes of visual system models [5, 6, 7]. Although several studies have found benefits of adding recurrence [8, 9, 10, 11], others report mixed results or found that recurrence does not serve the same function as in the brain [12, 13, 14]. These inconsistent findings may stem from the wide variety of architectural choices, including recurrence type, connectivity pattern, temporal formulation, that vary across studies without systematic comparison.

A key limitation of most existing recurrent convolutional neural networks (RCNNs) studies is their reliance on discrete-time recurrence, typically unrolled for only a handful of time steps. This coarse-grain approximation cannot capture the full complexity of continuous neural dynamics and is at odds with traditional methods in computational neuroscience, which treat neural circuits as dynamical systems governed by differential equations. Yet using continuous-time models in a machine learning setting poses engineering challenges: biologically realistic dynamics require small simulation step sizes (on the order of milliseconds), yielding tens to hundreds of time steps per stimulus and a correspondingly large computational graph during backpropagation through time. The implementation of such networks must therefore be efficient enough to make systematic scientific exploration feasible [15, 10].

Despite recent explorations of recurrent connections in biological vision models, fundamental questions remain unresolved about how to effectively model these dynamics. Can biologically plausible temporal dynamics emerge from simple recurrent connections combined with metabolic constraints, or are explicit normalization mechanisms necessary to replicate cortical response properties such as adaptation and sublinear temporal summation? Do different patterns of cortical connectivity such as dense lateral connections versus spatially organized feature-specific connections give rise to functionally distinct dynamics, or are they computationally equivalent? To what extent do different parameter choices across studies impact results? And must biological realism come at the cost of computational efficiency and task performance? These questions have proven difficult to address systematically because existing tools either sacrifice biological realism for computational efficiency or lack the flexibility to explore the parameter space comprehensively. Even state-of-the-art recurrent models fail to reproduce accurate within-area dynamics [16]. This gap is compounded by the limitations of current model comparison approaches, which rely on aggregate benchmark scores that can mask qualitative differences between models [17, 18]. Answering these questions requires an approach that enables controlled manipulation of architectural components while maintaining both biological plausibility and computational tractability.

To address these challenges, we introduce DynVision, a modular open-source toolbox for constructing and evaluating RCNNs with continuous-time dynamics. Inspired by recent work combining differential equations with deep convolutional networks [19, 20, 21], DynVision implements numerical ordinary differential equation (ODE) with heterogeneous delays, supports several biologically motivated recurrence types, and separates modeling decisions from implementation details through a configuration-driven design. We demonstrate the toolbox by systematically exploring how architectural choices like recurrence type, integration target, and temporal parameters shape learned dynamics, comparing model responses against cortical recordings, and revealing that different recurrent configurations give rise to functionally distinct regimes: one supporting temporal normalization, another enhancing noise robustness.

## 2 Methodology

Here, we provide a description of the overarching philosophy and core functionalities of the DynVision toolbox. For a more detailed account of all functionalities, consult the DynVision software documentation. Code can be found at: https://github.com/Lindsay-Lab/DynVision/.

### 2.1 Design Approach

On the one end of the neural network spectrum there are detailed models involving morphologies, cell-types, and spikes that focus on simulating realistic neural activity (e.g. those built with NEST [**?** ], Neuron [22], BRIAN [23]). On the other end, there are abstract network models often based on machine learning methodologies that focus on optimized behavioral performance, such as classification (e.g. VGG, ResNet, Transformers). Complementary to the modeling efforts on either extreme, “NeuroAI” models focus on information processing with biologically plausible components at a level of abstraction that enables both computational ability and meaningful comparison with neural and behavioral data [24]. This approach aligns with the broader view that recurrent neural networks serve as tractable surrogates for biological neural dynamics, inheriting the temporal and geometrical properties of the systems they model [25]. DynVision supports this modeling niche wherein biological realism and performance optimization are both required for the purposes of studying how biological features give rise to behavior.

Components of an artificial neural network can be mapped to biology in a variety of ways and with varying levels of specificity. Multi-layer CNNs, for example, parallel the hierarchical compositionality of the ventral visual stream (V1 -> V2 -> V4 -> IT), with early layers detecting simple features through small receptive fields analogous to V1 simple cells and additional pooling operations representing the function of complex cells [5]. Deeper layers progressively combine these features into increasingly abstract representations and larger receptive fields. Weight sharing and spatial pooling in CNN models realize a translational invariance that support the learning of generalizable visual features and the recognition of objects regardless of their precise position. Imitating the feedforward connections between pyramidal neuron populations of the cortical areas, the convolutional tensor operations maintain a spatial correspondence of the visual field, representing the retinotopic organization found in the visual cortex.

Here, we interpret each entry in the 3D activation tensor ([`n_channels, y_dim, x_dim`]) as a unit representative of a single neuron or a population of neurons, with the activation value corresponding to an average spiking rate. This interpretation establishes direct correspondences between network components and cortical structures, such as recurrent connections within cortical areas. Horizontal recurrent cortical connections are typically mediated by interneuron populations [1]. These recurrent synaptic connections are spatially organized favoring local connectivity [26]. Because recurrent convolutions combine local information from neighboring tensor units, the convolutional kernel realizes a local connectivity kernel. As detailed more in Section 2.3.1, different types of convolutional kernels can therefore model different biological connectivity patterns.

Similarly, skip and feedback connections model the long-range pyramidal neuron connections between cortical areas that integrate information from different hierarchical processing levels. For integrating skip, feedback, or recurrent activity into the feedforward processing of a layer, various strategies can be employed that parallel different biological integration mechanisms, such as additive or multiplicative modulation [21, 20]. These synaptic connection types have conduction delays that correlate with the spatial distance between neurons. Transmission delays in neural networks are realized by storing past activity in hidden states that can be accessed after a certain number of timesteps. As the hidden state buffers of our models hold several timesteps of activity; accessing different timestep slots in the hidden states can represent the different conduction delays for different connection types (see Section 2.2.2).

Nonlinearities, like ReLU, can be interpreted to represent the all-or-nothingness of action potential generation which only occurs when the membrane potential exceeds a threshold value (i.e. a bias), and the dynamical systems formulation approximates the aggregate membrane potential dynamics (see Section 2.2). With this interpretation of the architecture elements, recording the activations at different points in a layer’s order of operations corresponds to measuring the electrical activity at different points of the neuron’s cell body. Activations after a convolution would correspond to post-synaptic potentials in the dendrites, activations after the integration of recurrent, skip, or feedback signals and before the nonlinearity would correspond to the soma membrane potential, and activations after the nonlinearity would correspond to the generation of action potentials in the axon. The targeted recording of activations can further be used for an activity loss that mimics the metabolic costs of generating and transmitting electrical signals in the brain (see Section 2.4).

The biological interpretation of these components guides the design of the network architecture, including the order of operations within each layer, e.g convolution > bias > dynamics step > nonlinearity > pooling. As different interpretations about the components and the tensor unit activations impact the network architecture, e.g. where and how recurrent activity is integrated with each layer, our toolbox framework provides the flexibility to customize these design choices (see Section 2.3.4).

In the following sections we provide an overview of the options our toolbox provides to users, motivated by what is needed in order to efficiently explore biologically-inspired dynamics.

### 2.2 Temporal Dynamics

#### 2.2.1 Dynamical Systems Description

In biological neural networks, activity evolves over time and space in a structured and continuous manner based on past input and intrinsic network properties. DynVision implements numerical ODE solvers to build RCNNs with more precise and realistic temporal dynamics. This is often modeled within a dynamical system formulation as the differential equation in Figure 1 [20, 19]. The toolbox incorporates this dynamical systems formulation by accordingly evolving the activity over time using a stepwise numerical differential equation solver (Euler method, Figure 1 (2)), which evolves a layer’s activity on a timescale *τ* for time *t* based on the layer’s activity at previous time *t* − *dt* and any input to this layer arriving at that time. Thus, the step size *dt* determines the temporal resolution of the discrete approximation to the continuous system.

**Figure 1:**
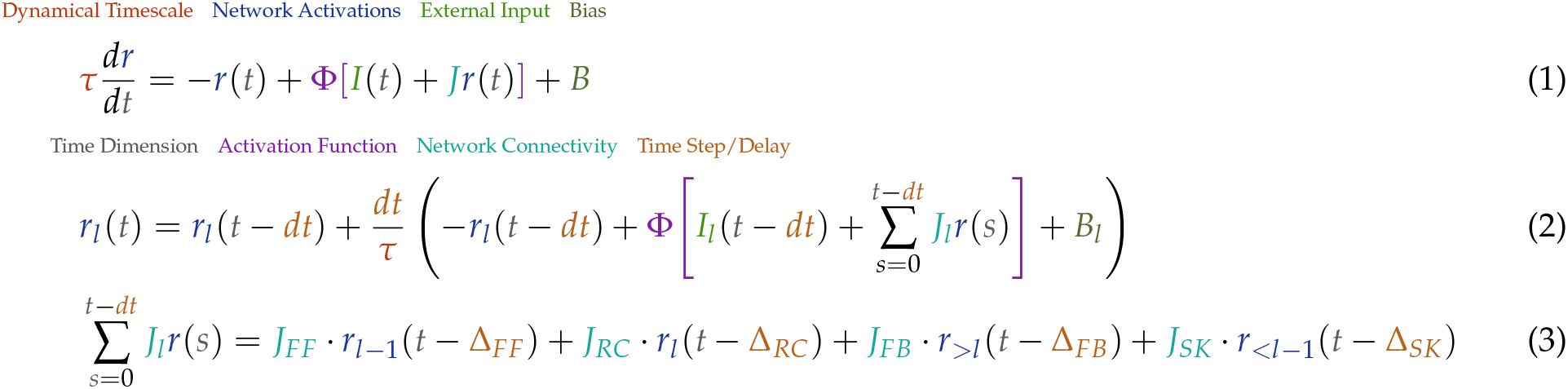
Dynamical systems formulation of the neural network activity with heterogenous connectivity and delays. **1)** Differential equation describing the evolution of network activity over time given the connectivity, and input, with a timescale given by *τ*. **2)** A numerical stepwise approach to approximately solving the differential equation via the Euler method for the activity *r*_*l*_ in layer *l* for the subsequent time step *t*. The contributions by feedforward (FF), lateral recurrent (RC), feedback (FB), and skip (SK) connections are expanded in equation **3**). Notably, delays Δ need to be integer multiples of *dt*.

Note that some details of this formalism can be modified by reordering the order of operations that are executed for each layer (with the `layer_operations` list parameter; see Section 2.3.4). This can include placing certain inputs outside of the activation function or moving the bias outside of the temporal evolution step.

#### 2.2.2 Heterogeneous Delays

Different neural connections are governed by different temporal delays, reflecting conduction velocities and the spatial distance between the neurons. Our toolbox allows for temporal unrolling with separate time delays for different types of connections between network layers. Specifically, at time *t* a layer can receive input from its own past activity *r*_*l*_ at *t* − Δ_*RC*_ via the laterally recurrent connections *J*_*RC*_, feedforward input from the preceding layer *r*_*l* −1_ at *t* − Δ_*FF*_ via the connectivity *J*_*FF*_, and recurrent feedback connections from a succeeding layer *r*_>*l*_ via *J*_*FB*_ at *t* − Δ_*FB*_. By varying these parameters, users can systematically evaluate the influence of different temporal delays on network dynamics.

#### 2.2.3 Unrolling of Time

This system of heterogeneous delays can be naturally applied when unrolling the temporal graph in “biological” time, wherein activity propagates through the depth of the network over timesteps. But under many circumstances it can just as well be used when unrolling in “engineering” time, wherein activity propagates from the input to output layer within a single timestep. By simply setting the feedforward delay 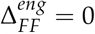 we effectively treat the network as an instantaneous system, akin to unrolling in engineering time (Figure 2). To switch from biological to engineering time in a network with skip and feedback connections, the feedforward delay needs to be subtracted from the skip delay Δ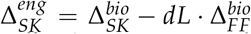 with *dL* being the number of layers spanned by the skip connection; while for feedback connections the feedforward delay needs to be added to the feedback delay 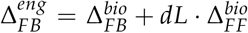. The use of engineering time only works while 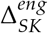 is positive; thus engineering time requires that skip connections are equal or faster than multi-synaptic feedforward pathways.

**Figure 2:**
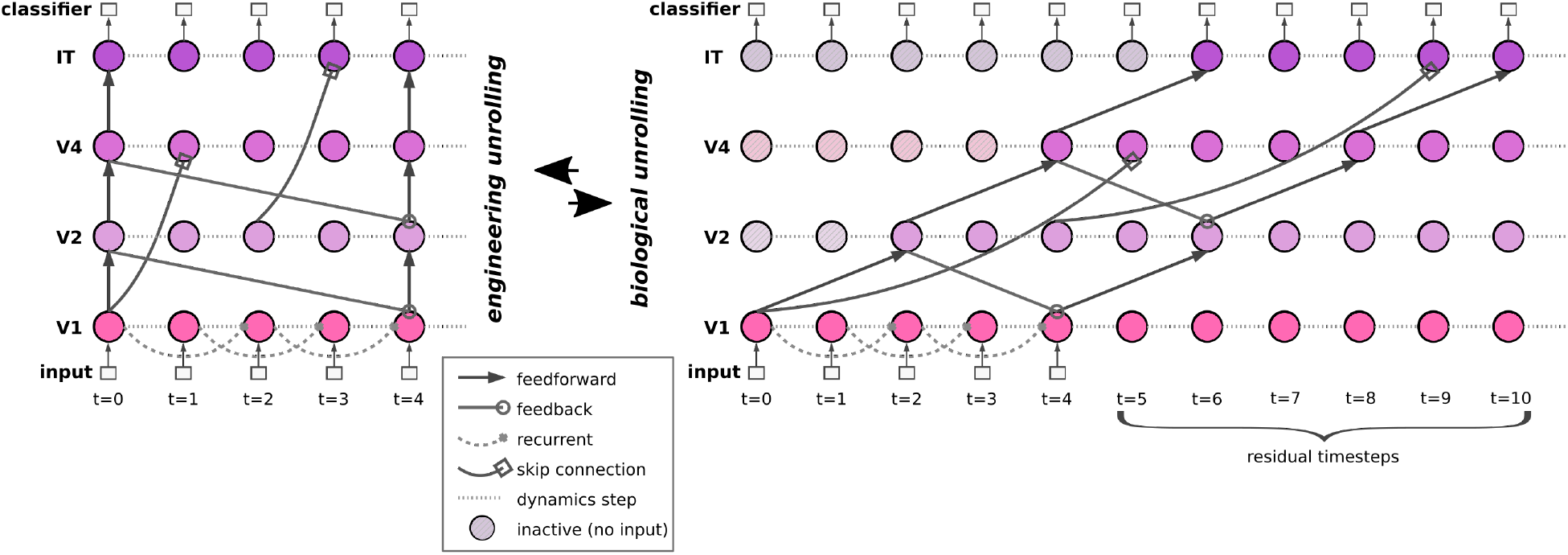
Network with recurrent, skip, and feedback connections unrolled in engineering versus biological time. In biological time (*right side*), the network has feedforward connections with delay Δ_*FF*_ = 2, recurrent connections with Δ_*RC*_ = 2, skip connections over two layers with Δ_*SK*_ = 5, and feedback connections over one layer with Δ_*FB*_ = 2. Note that each node has equivalent in- and out-put connections, but only a few are shown here for illustration purposes. By unrolling the network instead in engineering time (*left side*), the delays become Δ_*FF*_ = 0, Δ_*RC*_ = 2, Δ_*SK*_ = 1, Δ_*FB*_ = 4, but the signal flow through the network and time remains identical.

In practice, these delay transformations may need to be adjusted by ± 1 depending on the order in which feedforward delay, skip, and feedback integration are applied. For most architectures the skip delay index needs to be incremented since the layer computations of its source for that timestep were already executed, while source layer computations for a feedback input for the same timestep are not.

To make sure an input with a given number of timesteps is completely processed by the classifier, the simulation time is automatically extended by the number of residual timesteps it takes for the first input signal to reach the final layer. When simulating the network in engineering time, the residual timesteps are per definition zero.

Simulating a model in engineering time is more computationally efficient. In an example 3.5h training run using our DyRCNNx8 model with full recurrence (Section 3.1) on CIFAR-100 with 30 timesteps (60 ms) we use the following delay values for engineering and biological time: Δ_*FF*_ = (0 | 10) ms, Δ_*RC*_ = 6 ms, Δ_*SK*_ = (2 | 22) ms, Δ_*FB*_ = (30 | 10) ms. The training speed as time per epoch decreases by ∼ 29% and memory demand decreased from 2.39 GB to 2.13 GB for unrolling in engineering time compared to biological time. Note: engineering time reduces the number of simulation steps but it can increase the length of the hidden state due to larger Δ_*FB*_, which can lead to higher memory demands depending on the settings.

The recurrence delay (Δ_*RC*_ = 6 ms) and time constant (*τ* = 5 ms) are independent of the type of unrolling and chosen to approximate biologically observed timescales and scale proportionally with the temporal resolution *dt*, while the feedforward delay is a free parameter determined only when converting between engineering and biological time conventions.

#### 2.2.4 Temporally Varied Input Presentation

When modeling temporal model dynamics, the input has to be presented over multiple timesteps. The input can be presented as a static image over time, combined with null inputs for certain timesteps, or potentially undergo more complex time-dependent transformations (e.g., by varying the contrast or by showing dynamic stimuli).

The toolbox provides three complementary approaches for extending static images over time, each suited to different experimental contexts. For high-performance training with FFCV (Fast Forward Computer Vision) dataloaders [27], setting `data _timesteps> 1` creates temporally extended sequences during the data loading phase, minimizing the CPU-GPU transfer overhead. For flexible testing and experimentation, setting `n_timesteps > 1` in the model enables dynamic temporal expansion during the forward pass. Alternatively, specialized PyTorch dataloaders can implement complex temporal transformations for specific experimental protocols. These options provide flexibility in balancing computational efficiency, biological realism, and experimental control depending on the research question.

The time extension at runtime can be varied by setting `data_presentation_pattern` to any boolean string such as `01110`, indicating a temporal sequence of null input (`0`) and input image (`1`). The pattern string is always proportionally stretched or compressed internally so that its length equals `n_timesteps`, and thus each value corresponds to one timestep. The same functionality is also available by using the time extension with PyTorch dataloaders with the option to also realize more complex temporal transformations.

Overwriting and parameterizing the iterator of the standard PyTorch dataloader allows for different testing scenarios analogous to common experimental protocols. We provide several of these out of the box. For example, our `StimulusDuration` loader presents an image after a delay *δ*_*i*_ for *δ*_*s*_ duration followed by silent outro period *δ*_*o*_; the `StimulusInterval` behaves similarly besides also presenting a second identical stimulus with a *δ*_*d*_ delay after the first; the `StimulusContrast` loader adds to the StimulusDuration setup by multiplying the image input with a scalar to vary its contrast.

#### 2.2.5 Stimulus Noise

To investigate model robustness to degraded sensory input DynVision provides a `StimulusNoise` dataloader that adds controlled noise to stimuli across timesteps. The implementation supports multiple noise types (Gaussian, uniform, salt-and-pepper, Poisson) with configurable signal-to-signal-plus-noise (SSNR) ratios, enabling systematic investigation of noise robustness under controlled conditions. Users can specify whether the noise pattern is held static across timesteps, drawn independently per timestep, or temporally correlated, allowing investigation of how the temporal structure of noise interacts with recurrent dynamics. The noise parameterization is fully configurable through the YAML configuration files, ensuring reproducibility and enabling systematic exploration of how different recurrent architectures and training regimes affect robustness to various forms of input degradation.

#### 2.2.6 Idle Timesteps

Initializing hidden states in recurrent models is realized in various ways. CORnet initializes hidden states with zeros, whereas the hidden states of our DyRCNNx8 model produce `None` (i.e. no recurrent signal integration) when they are not yet populated. CordsNet addresses this issue by preceding its forward pass with a number of idle timesteps, where only a null input is provided, to allow self-generated activity to stabilize and populate the hidden states. In DynVision this is implemented with a memory-efficient idle timestep mechanism using a cache-reset-restore pattern: the network processes null input (by default zero tensors representing neutral, contrast-free input at the normalized mean pixel value) for a specified number of idle timesteps to allow spontaneous activity to converge, then caches the resulting hidden states, resets the computational graph, and initializes the stimulus processing with these converged states. This approach enables pre-stimulus convergence without gradient flow complications or memory accumulation during the idle period.

### 2.3 Network Connectivity

#### 2.3.1 Lateral Recurrent Connection Types

DynVision contains several options for recurrent connectivity patterns. These options represent different beliefs about the spatial and feature extent of lateral connections. Specifically:

**Self-recurrence** connects a unit only to itself. Input is determined by multiplying the layer’s activation tensor with a scalar weight ([8]), mimicking connections within a local group of neurons (Figure 3).

**Figure 3:**
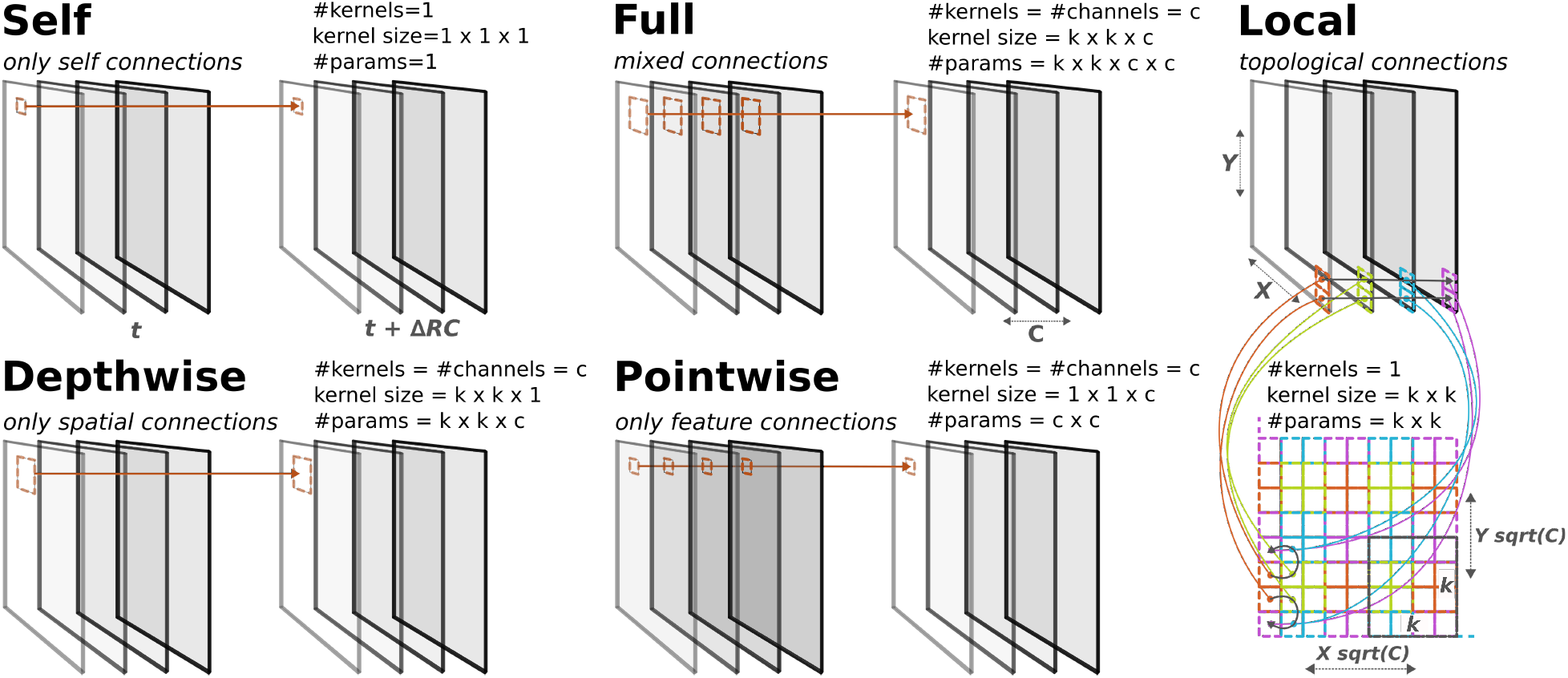
Five basic types of kernel convolutions used to realize recurrent connections. Each convolution type is represented by its input 3D tensor (Height x Width x Channels) at time *t* and its output 3D tensor at time *t* + Δ_*RC*_ which is to be integrated with the feedfoward activations. The convolutions differ in their kernel dimensions and number of individual kernels (Height, Width, Channels)x(# of Kernels) that can be parameteric with the kernel scale *k* and the number of channels *c*: Self (1,1,1)x(1), Full (k, k, c)c(c), Depthwise (k,k,1)x(c), Pointwise (1,1,c)x(c), Local (k,k)x(1). The Local recurrence introduced here involves a mapping from the 3D tensor to a 2D “cortical” plane on which the convolution is applied.

**Full recurrence** applies a standard kernel convolution so that a unit is influenced by a nearby spatial region across all channels, capturing broad lateral connectivity across feature maps. This was introduced by [28] and is frequently used in the literature (e.g. [29], Figure 3).

**Depthpointwise recurrence** instead of a full convolution, uses a depthwise (spatial dimension) and then a pointwise (feature dimension) convolution (see Figure 3). This is also known as a depthwise separable convolution and has been used by the computer vision community ([30]). While not commonly used in neuroscientific models, it does map onto the structure of lateral recurrence in the visual system as observed, for example, in orientation pinwheels and topographic maps. Specifically, visual neurons tend to get input from other neurons that 1.) represent the same spatial location but have different feature preferences (depthwise) and 2.) have the same preferred features but represent different locations in space (pointwise) [31]. Depthwise separable convolutions are desirable from an engineering perspective as they reduce the number of parameters compared to a full convolution.

**Pointdepthwise recurrence** here, compared to the depthpointwise recurrence, the order of the two partial convolutions is inverted. This follows from the idea that neurons representing similar locations in space are arranged closer together in cortical space so that a signal propagation across features (pointwise) would precede the signal exchange across locations (depthwise) (Figure 3).

**Local recurrence** aims to capture the 2-D topology of the cortical sheet. Here, all units in a layer are systematically arranged on a 2-D grid (inspired by cortical organization such as orientation pinwheels in V1 ([26], see Figure 3). To efficiently precompute this mapping, the module initialization requires the shape information of the input tensor (`dim_y` and `dim_x`). A convolution with kernel size > 1 is applied to this grid. The result is that input to each unit is a combination of cortically-local feature and space information (Figure 3). Consistent with this, connectivity-constrained recurrent models that penalize long-range wiring were shown to develop domain-selective topographic organization resembling primate inferotemporal cortex [32]. This approach is inspired by LLCNNs [33, 34] but also incorporates spatial retinotopy in the 2-D map. Our approach has similarities to other topographic mappings like All-TNN [35] and TDANN [36], which also map to a combined feature-spatial plane, but have an explicit spatial loss instead of doing a convolution with weight-sharing on this plane. This mapping to a 2-D topology requires that the number of channels is a square. If this is not the case, the mapping extends the number of channels to the next square number by duplicating the last channels as with a reflecting boundary. The supplemented channels are removed again in the mapping back to the 3-D tensor.

**Local depthwise recurrence** The local recurrence can be extended with an additional depthwise convolution mimicking patchy long-range connections in the visual cortex [37] between units with neighboring receptive fields but the same feature preferences. Typically, connectivity matrices *J* are implemented via kernel convolutions, reducing the number of parameters to learn (biologically, this corresponds to an assumption that local connectivity statistics are similar for neurons across the cortical sheet).

In our toolbox, the module `RecurrentConnectedConv2d` combines a feedforward convolution (`torch.nn.Conv2d`) operation with a recurrence operation of choice (`recurrence_type`) and handles the storage and integration of corresponding hidden states with a variable time delay (`t_recurrence`). See Section 2.3.3 for how (recurrent) signals are integrated. The module can be extended with a second feedforward convolution by setting a `mid_channels` attributes, as in several popular architectures [15].

Recurrent connections are trained with a reduced learning rate, by default 5x smaller than feedforward connections (all hyperparameters described in Table 1). This design choice is motivated by biological evidence that recurrent synaptic plasticity tends to be weaker and operate on slower timescales than feedforward plasticity [38, 39], and helps stabilize training dynamics in continuous-time networks where recurrent connections can otherwise lead to runaway dynamics.

**Table 1:** Default Training Configuration.

| Parameter | Value | Description |
| --- | --- | --- |
| **Temporal Dynamics** |  |  |
| Time steps | 20 | Number of simulation timesteps |
| Resolution ($dt$) | 2 ms | Integration time step |
| Time constant ($\tau$) | 5 ms | Neural time constant |
| Feedforward delay ($\Delta_{FF}$) | 0 ms | Engineering time |
| Skip connection delay ($\Delta_{SK}$) | 0 ms | Engineering time |
| Recurrent delay ($\Delta_{RC}$) | 6 ms | Recurrent connection delay |
| Integration strategy | Additive | Activity integration method |
| Dynamics solver | Euler | Numerical ODE solver |
| Recurrence target | Output | Recurrence applied to layer output |
| **Architecture Configuration** |  |  |
| Skip connections | Enabled | V1→ V4, V2→ IT |
| Feedback connections | Disabled | V1← V4, V2← IT |
| Normalization | None | No BatchNorm/LayerNorm |
| **Training Setup** |  |  |
| Training epochs | 200 | Total training duration |
| Datasets | ImageNette | 224px, 10 classes |
| Batch size | 192 | Training batch size |
| Initial learning rate | 0.0008 | Base learning rate |
| LR scheduler | CosineAnnealingLR | $T_{max} = 500$ |
| Recurrent LR factor | 0.2 | 5x smaller LR for recurrent weights |
| Feedback LR factor | 0.5 | 2x smaller LR for feedback weights |
| Optimizer | Adam | Optimization algorithm |
| Weight decay | 0.0005 | L2 regularization |
| **Weight Initialization** |  |  |
| Feedforward weights V1 | Kaiming | $\mu = 0, \sigma = 0.053$ |
| Feedforward weights V2 | Kaiming | $\mu = 0, \sigma = 0.050$ |
| Feedforward weights V4 | Kaiming | $\mu = 0, \sigma = 0.035$ |
| Feedforward weights IT | Kaiming | $\mu = 0, \sigma = 0.025$ |
| Recurrent weights | Truncated Normal | $\mu = 0, \sigma = 0.004$ |
| Skip & Feedback weights V1↔V4 | Truncated Normal | $\mu = 0, \sigma = 0.001$ |
| Skip & Feedback weights V2↔IT | Truncated Normal | $\mu = 0, \sigma = 0.001$ |
| Bias initialization | Zero | All biases set to 0 |
| **Loss Function** |  |  |
| Primary loss | Cross-entropy | Classification loss |
| Auxiliary loss | Activity Loss (x0.0002) | Biological constraint |
| Loss reaction time | 4 ms | Temporal offset for loss calculation |
| **Optimization** |  |  |
| Gradient accumulation | 4 batches | Effective batch size: 768 |
| Precision | bf16-mixed | Mixed precision training |
| Gradient clipping | Norm, value=1.0 | Gradient norm clipping |
| **Hardware** |  |  |
| GPU | NVIDIA A100 | Single GPU core |
| CPU | 16 cores | Parallel data processing |

#### 2.3.2 Skip and Feedback Connections

Besides lateral recurrent connections that link units within one layer across time steps, skip and feedback connections link units across layers and time steps. Anatomical studies reveal that feedback projections from higher cortical areas like V4 and IT back to V2 and V1 are as numerous as feedforward connections, suggesting a fundamental role in visual computation [3, 2].

Skip and feedback connections describe a similar process: a copy of a layer’s output is diverted from the immediate feedforward path to be integrated with a downstream (skip) or upstream (feedback) region. Thus, we provide a generalized module implementation that covers both cases, and use the same integration strategies as available for the recurrent connection (e.g. additive or multiplicative). The shapes of the source and target layers may not necessarily match, in which case an up or downsampling operation adjusts the spatial dimensions.

To avoid requiring the user to determine the right layer shapes and corresponding transformations in advance, the toolbox offers an `auto_adapt` option, so that the connection can be defined by referencing a source and target layer, and the correct transformation is created upon the first forward pass. This facilitates architecture exploration because connections between layers and orders of operation can easily be swapped without causing errors.

#### 2.3.3 Integration of (Recurrent) Connections

Some studies suggest that recurrence may be best modeled as a gain modulation [40, 41]. Therefore, in DynVision, the way the recurrent signal is integrated with bottom-up input can be set by `integration_strategy` to either `additive` (*x*′ = *x* + *h*) or `multiplicative` (*x*′ = *x* ∗ (1 + *torch*.*tanh*(*h*))) or a custom callable function. To explore the impact of recurrent connections, during inference the recurrence can be disabled by setting `feedforward_only = True`.

For the recurrence, the layer output (extracted at the `delay` operation, see Figure 4) is convolved according to the recurrence type and the result is integrated with the feedforward activation tensor. The location at which it is integrated with respect to the feedforward convolutions can be selected with the argument `recurrence_target`:

`output|` : Integration with the activation tensor after the last layer convolution and before the final nonlinearity. Integrating after the convolution(s) allows direct matching of tensor dimensions, and is therefore a popular choice in the literature [e.g. in 19].

`input|` : Integration with the layer input before the first convolution (see example in 4).

`middle|` : *Only applicable if the layer consists of two convolution operations*. The recurrence output is integrated immediately before the second convolution, inspired by the CORnet architecture [15]).

In case there is a mismatch in the spatial dimensions (for the `input` or `middle` recurrence target) of the recurrent activity tensor with the feedforward activity tensor, an additional up or downsampling (using nearest neighbors) after the recurrence convolution.

#### 2.3.4 Order of Layer Operations

The model base classes provide a structure to name and execute the operations of each layer in a flexible order. The activations of each layer are computed by calling its layer operations in the order defined by the `layer_operations` attribute. If a given operation is not defined for a layer, it is skipped.

For example, the order of operations used in the DyRCNN models presented in this paper is (unless otherwise noted):

~~~
layer_operations:
-“rconv” # apply recurrent convolutional module
-“addskip” # add activity from skip connections
-“addfeedback” # add activity from feedback connections
-“tstep” # apply dynamical systems ode solver step
-“nonlin” # apply nonlinearity
-“record” # record activations in storage buffer
-“delay” # set and get delayed activations
-“pool” # apply pooling
~~~

**Figure 4:**
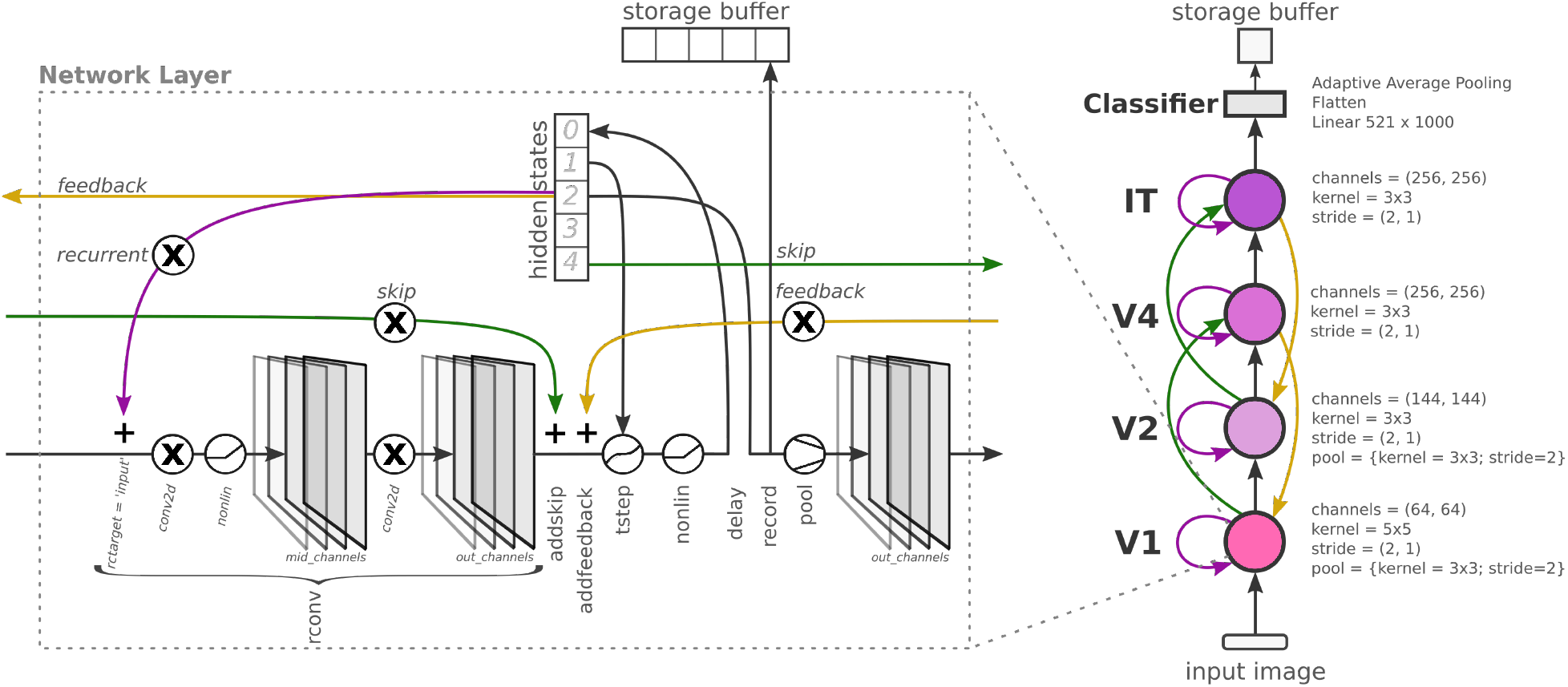
Architecture schematic of the DyRCNN model family, native to the DynVision toolbox. The left hand side shows the signal flow and processing within one network layer. The layer operations are shown as circular pictograms or the merging or splitting of arrows, i.e., ‘+’ symbolizes a piecewise addition of the tensors. The layer units are represented by their 3D activation tensors. The order of layer operations (vertical labels) can be flexibly arranged as described in Section 2.3.4. Recurrent activity can be integrated with the `input` (shown here), `middle`, or `output` activations of the convolutions. Pointers to specific timestep slots of the hidden state indicate the respective delays (Section 2.2.2), here Δ_*FF*_ = 2, Δ_*RC*_ = 2, Δ_*FB*_ = 2, Δ_*SK*_ = 4. The right hand side shows our four-layer network, the layer parameters, and the connections between layers. Note that the pooling operation is only applied in V1 and V2. The tuple parameters indicate the individual values for the two convolutions in the respective layer, when they differ from each other.

The naming of the layers and operations is chosen by the user, except for the reserved operation names: `delay`, which writes the current activity into the hidden states and retrieves past activity with the correct delay; `tstep`, which accesses the most recent hidden state); and `record`, which stores the current activity in the model’s storage buffer.

Any custom model only needs to define its layer names (e.g. `self.layer_names = [‘V1’, ‘V2’, ‘V4’, ‘IT’]`) and modules in `_define_architecture()`, following the naming convention `self.<operation>_<layer_name> = <module>`. For layer-unspecific operations, e.g. a nonlinearity, the `_<layer_name>` can be omitted.

### 2.4 Loss Functions

The toolbox supports the application of multiple loss functions to be selected in the configs. Any loss function-specific configurations are set in the configs as dicts with `loss_config: [<loss_name>: <loss_configs>]`. A weighting of the individual loss contributions is coordinated with a `weight` scalar value in the loss configs. The available loss functions can be extended by adding a corresponding implementation or wrapper class for a pytorch loss function to the `dynvision/losses/` folder. The BaseLoss parent class provides consistent input processing, shape validation, and reduction over the batch and time dimension.

The default configuration is `loss_config: [CrossEntropyLoss: [weight: 1], ActivityLoss: [weight: 0.0002]]`. These represent task accuracy and neural activity sparseness as follows:

#### Category Loss

Per default we use the widely used cross-entropy-loss function as a category loss. The category loss automatically ignores any model output for the initial number of residual timesteps, for which model input hasn’t yet reached the classifier. Additionally, there is a `loss_reaction_time` setting that further removes the initial *n* timesteps (after any residual timesteps) from consideration for the category loss as desired.

#### Activity Loss

Activity regularization promotes more stable network activity and is biologically motivated by mechanisms of homeostatic plasticity [42]. In the toolbox, we add an effective activity regularization via an activity loss function (inspired by [43]) that penalizes large unit responses, thus mimicking the metabolic constraints of action potential generation in the brain. This loss function accesses the activations after a defined selection module (depending on mode) from the computational graph without the need for any additional storing of activation tensors. The activity loss is defined as the p-norm (default is *p* = 1) of the activation tensor *r*_*m*_ of module *m* divided by its number of units *N*_*m*_ and averaged across all modules *M* (Eq. 4).

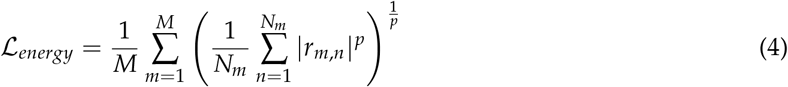

The activity loss function implementation has several presets determining which activations in the layer’s order of operation is used for the loss. `PSP`: post-synaptic potentials, i.e. activations after convolutional operators; `AP`: action-potentials, i.e. activations after nonlinearities; `EI`: excitatory-inhibitory balance (default), i.e. just before nonlinearities, effectively incentivizing a balance between negative and positive input values.

Adding a weight decay (default 0.0005) further adds weight regularization to promote network stability and more efficient and sparse coding, further modeling metabolic constraints.

### 2.5 Reference Models

To facilitate rapid experimentation and benchmarking, the toolbox integrates implementations of several well-established neural network architectures from the literature, including AlexNet [44], the ResNet family [45], CORnet-RT [46], and CordsNet [19]. The latter two also integrate types of recurrent connections which we explore in detail in Section 3.11. In addition to these models, users can easily instantiate the showcase models introduced in this paper, DyRCNNx8, and its downscaled versions DyRCNNx4 and DyRCNNx2. The framework also supports the automatic loading of pre-trained weights, with mechanisms in-place to correctly match parameters even when model configurations have been modified (e.g., additional layers or biologically inspired features added). Notably, models are designed to be data context-aware, meaning they are automatically initialized with the appropriate number of output classes based on the provided dataset. The toolbox further enables selective training of only non-pre-trained parameters or full model fine-tuning, offering flexibility for transfer learning applications.

### 2.6 Software Design

DynVision is designed with scientific exploration in mind. We follow best principles for scientific software engineering to make the toolbox transparent and easy to use.

#### Interoperability

To integrate technical advances without reinventing infrastructure, the toolbox builds on: PyTorch [47] for performant tensor operations; PyTorch Lightning [48] for training orchestration and hardware allocation; FFCV dataloaders [27] to accelerate the data loading bottleneck; Snakemake [49] for automated, scalable workflow management; and YAML with Pydantic^1^ for human-readable, type-safe parameter interfaces.

#### Modular Components

The fundamental advantage of modeling approaches in neuroscience is that, contrary to actual brains, models allow for precise control and monitoring of every variable. DynVision enables this through complementary modularity mechanisms. First, each model is defined as a sequence of layers and each layer as a sequence of operations, so that components such as recurrence types, dynamics solvers, and signal delays can be reordered and swapped without changing overall model handling. Second, core capabilities are implemented as five specialized base classes combined via multiple inheritance: `TemporalBase` manages the time dimension and executes layer operations in configurable order; `LightningBase` handles mixed-precision training, checkpointing, and multi-GPU scaling; `StorageBuffer` selectively records activations and hidden states across timesteps for comparison with neural data; `DtypeDeviceCoordinator` ensures dtype and device consistency across components; and `Monitoring` provides structured logging with automatic detection of numerical issues such as NaN gradients. Together, these mechanisms let researchers run controlled experiments where specific components are varied while others are held constant (Supplementary Section 8.1.1).

#### Configuration-driven Modeling

DynVision separates scientific modeling decisions from implementation details through a YAML-based configuration schema. The advantages are i) a lower coding barrier for developing and testing complex models, ii) reproducibility and systematic parameter-space exploration, iii) making architectural choices (e.g. the order of layer operations or type of time-unrolling) explicit and visible, and iv) integrating technical advances independently from model design. Every experimental output is accompanied by a fully resolved configuration file recording provenance metadata for each parameter, ensuring complete reproducibility (Supplementary Section 8.1.2).

#### Data Handling and Workflow Management

A symbolic link-based data pipeline eliminates storage redundancy when partitioning datasets into subsets or defining custom class groupings, with automatic label-index remapping so that models trained on full datasets can be seamlessly evaluated on subsets. For temporal models that process image sequences over many timesteps, the toolbox provides efficient options to unroll time either during dataloading for more flexible dynamic input presentations or during runtime for faster processing on static images (Supplementary Section 8.1.3). The full model lifecycle from data acquisition through training, analysis, and visualization is orchestrated by Snakemake [49], which manages dependencies, parallelizes independent jobs, and scales from laptops to HPC clusters (Supplementary Section 8.1.4).

### 2.7 Evaluation Metrics

The toolbox implements evaluation metrics that generalize to temporally extended inputs and are automatically computed and aggregated across models and testing experiments within the workflow.

#### Accuracy

Classification accuracy is computed as the binary correctness indicator for each sample and timestep. For a batch of *N* samples with classifier readouts *c*_*i*_(*t*) and ground truth labels *y*_*n*_ at timestep 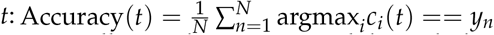. During training, accuracy is computed only for timesteps where non-null input has propagated through the network to reach the classifier, excluding residual timesteps and any loss reaction time offset. During testing, accuracy is computed for all timesteps, enabling analysis of classification dynamics during both stimulus presentation and subsequent null-input periods. Results are aggregated across samples, seeds, and stimulus categories according to experimental configurations. Top-*k* accuracy follows analogously, evaluating whether the correct label appears among the *k* highest-confidence predictions.

#### Confidence

Confidence measures quantify the model’s certainty in its predictions across time. Target confidence reports the classifiers softmax probability assigned to the correct class: 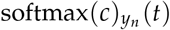, where *y*_*n*_ is the ground truth label. Guess confidence reports the maximum softmax probability: max_*i*_ softmax(*c*)_*i*_(*t*), corresponding to the model’s prediction. These metrics are particularly informative when combined with accuracy to characterize the temporal evolution of model certainty during stimulus processing and null-input periods.

#### Average Response

Layer-wise population activity is recorded by placing a `record` operation in the sequence of layer operations (Section 2.3.4), which stores unit activations via the `StorageBuffer` component (Supplementary Section 8.1.1). By default, activations are recorded after each layer’s nonlinearity. We further process the recorded unit activations by averaging them over each layer. This spatially aggregated signal provides a first-order proxy for comparison with experimental data such as electrocorticography (ECoG) recordings, which measure spatially averaged extracellular potentials.

## 3 Results

We built this toolbox to generalize biological RCNN network models and make them flexibly editable for systematic exploration. To demonstrate its capabilities, we focus on our primary working model, the DyRCNNx8, inspired by existing architectures in the literature. More so than feedforward networks, RCNNs present researchers with a large space of possible hyperparameter settings. Thus, we systematically explore the modeling parameter space to demonstrate RCNN task performance, identify optimal configurations, benchmark the effect of modeling choices, and then validate the biological plausibility of the resulting models by comparing against neural data and behavioral observations. Having established that our DyRCNN models can exhibit biologically realistic dynamics, we demonstrate the framework’s generality by showing it can precisely recreate established models from the literature (CORnet-RT and CordsNet).

All results presented in this work are based on models trained with at least three different random initialization seeds, with each seed producing different initial weight configurations. Throughout the Results section, unless otherwise noted, all plots showing average response activities include standard deviation errors as shading (often too small to see) computed across seeds and input samples.

### 3.1 Model Architecture

The version of the DyRCNNx8 model analyzed throughout this section contains variants of lateral recurrence and skip connections. We also briefly explore the impacts of feedback connections (more thorough explorations are reserved for future work). Notably, this model does not include any explicit normalization operation, to evaluate whether recurrence alone can realize biologically plausible solutions of normalization.

The architecture and order of operations is defined as follows:

**[V1] Conv2d**(*in=3, out=64, k=5×5, s=2*) **→ ReLU → Conv2d**(*in=64, out=64, k=5×5, s=1*) **→ +Recurrence** (Δ_*RC*_ = 6*ms*) **→ Euler Step** (*dt* = 2*ms, τ* = 9*ms*) **→ ReLU → Delay** (Δ_*FF*_ = 0*ms*) **→ MaxPool**(*3×3, s=2*)

**[V2] Conv2d**(*in=64, out=144, k=3×3, s=2*) **→ ReLU → Conv2d**(*in=144, out=144, k=3×3, s=1*) **→ +Recurrence** (Δ_*RC*_ = 6*ms*) **→ Euler Step** (*dt* = 2*ms, τ* = 9*ms*) **→ ReLU → Delay** (Δ_*FF*_ = 0*ms*) **→ MaxPool**(*3×3, s=2*)

**[V4] Conv2d**(*in=144, out=256, k=3×3, s=2*) **→ ReLU → Conv2d**(*in=256, out=256, k=3×3, s=1*) **→ +Recurrence** (Δ_*RC*_ = 6*ms*) **→ +Skip**(*V1*, Δ_*SK*_ = 0*ms*) **→ Euler Step** (*dt* = 2*ms, τ* = 9*ms*) **→ ReLU → Delay** (Δ_*FF*_ = 0*ms*)

**[IT] Conv2d**(*in=256, out=529, k=3×3, s=2*) **→ ReLU → Conv2d**(*in=529, out=529, k=3×3, s=1*) **→ +Recurrence** (Δ_*RC*_ = 6*ms*) **→ +Skip**(*V2*, Δ_*SK*_ = 0*ms*) **→ Euler Step** (*dt* = 2*ms, τ* = 9*ms*) **→ ReLU → Delay** (Δ_*FF*_ = 0*ms*)

**[Classifier] AdaptiveAvgPool2d**(*1*) **→ Flatten → Linear**(*529* → *10*)

Further parameters and training details can be seen in Table 1.

### 3.2 Equivalence of Engineering and Biological Time Unrolling

A useful feature of DynVision is the ability to simulate models using either engineering time (where Δ_*FF*_ = 0) or biological time (where Δ_*FF*_ > 0) unrolling. As described in Section 2.2, these two approaches produce mathematically equivalent dynamics when delays are properly adjusted. Contrary to previous work [10], this also works for models with top-down feedback connections. We validate this prediction by training a model in engineering time and testing it in both engineering and biological time configurations.

As shown in Figure 5, the model produces identical temporal dynamics (shifted by Δ_*FF*_) regardless of the unrolling scheme, confirming that researchers can choose the computationally more efficient engineering time for training while maintaining the ability to interpret results in biological time. This flexibility is particularly valuable when comparing model predictions to neural recordings where real-world propagation delays matter. In the following sections, all results are presented in engineering time so that the response onset of all layers is aligned at 0 for visual compactness.

**Figure 5:**
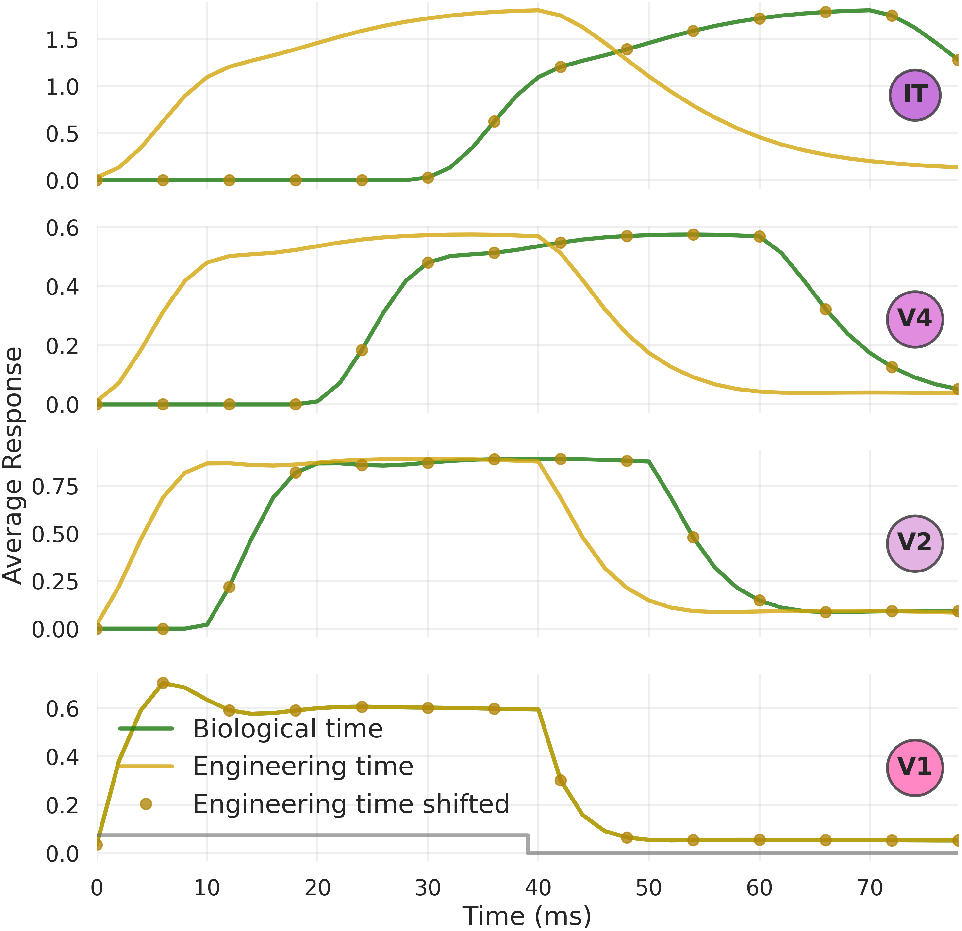
Equivalence of engineering and biological time unrolling. This DyRCNNx8 model was trained with full recurrence, skip (V1 → V4; V2 → IT), and feedback connections (IT → V2; V4 → V1) on input with 40 timesteps unrolled in engineering time (Δ_*FF*_ = 0 ms, Δ_*SK*_ = 0 ms, Δ_*FB*_ = 34 ms, Δ_*RC*_ = 6 ms, *τ* = 9 ms, *dt* = 2 ms), and then tested in both engineering time and biological time (Δ_*FF*_ = 10 ms, Δ_*SK*_ = 20 ms, Δ_*FB*_ = 14 ms, rest identical) on a single image presented for 35 timesteps followed by 35 timesteps of null input (indicated by the grey step function). Each configuration yields equivalent model responses, validating the mathematical framework and demonstrating computational flexibility.

### 3.3 Training Protocol: Activity Loss and Temporal Presentation Pattern

A critical consideration when training biologically plausible recurrent networks is balancing task performance with biological constraints. The activity loss function (Eq. 4) in `EI` mode (incentivizing excitatory-inhibitory balance) implements a metabolic constraint that promotes sparse neural coding. However, the relative weighting between the cross-entropy classification loss and the activity regularization loss affects both training dynamics and the final network behavior.

To establish appropriate training parameters for subsequent experiments, we systematically explored different hyperparameter setups. Figure 6 shows training and validation accuracy curves (panels A), as well as the corresponding evolution of the activity loss (panels B) and cross-entropy loss (panels C) for different relative weighting values of the activity loss component. Figure 6B show that the activity loss converges to a stable level relatively quickly, typically within 50-100 epochs, representing an efficient working point of the network. Once this regime is established, the cross-entropy classification loss continues to decrease as the model fine-tunes its representations.

**Figure 6:**
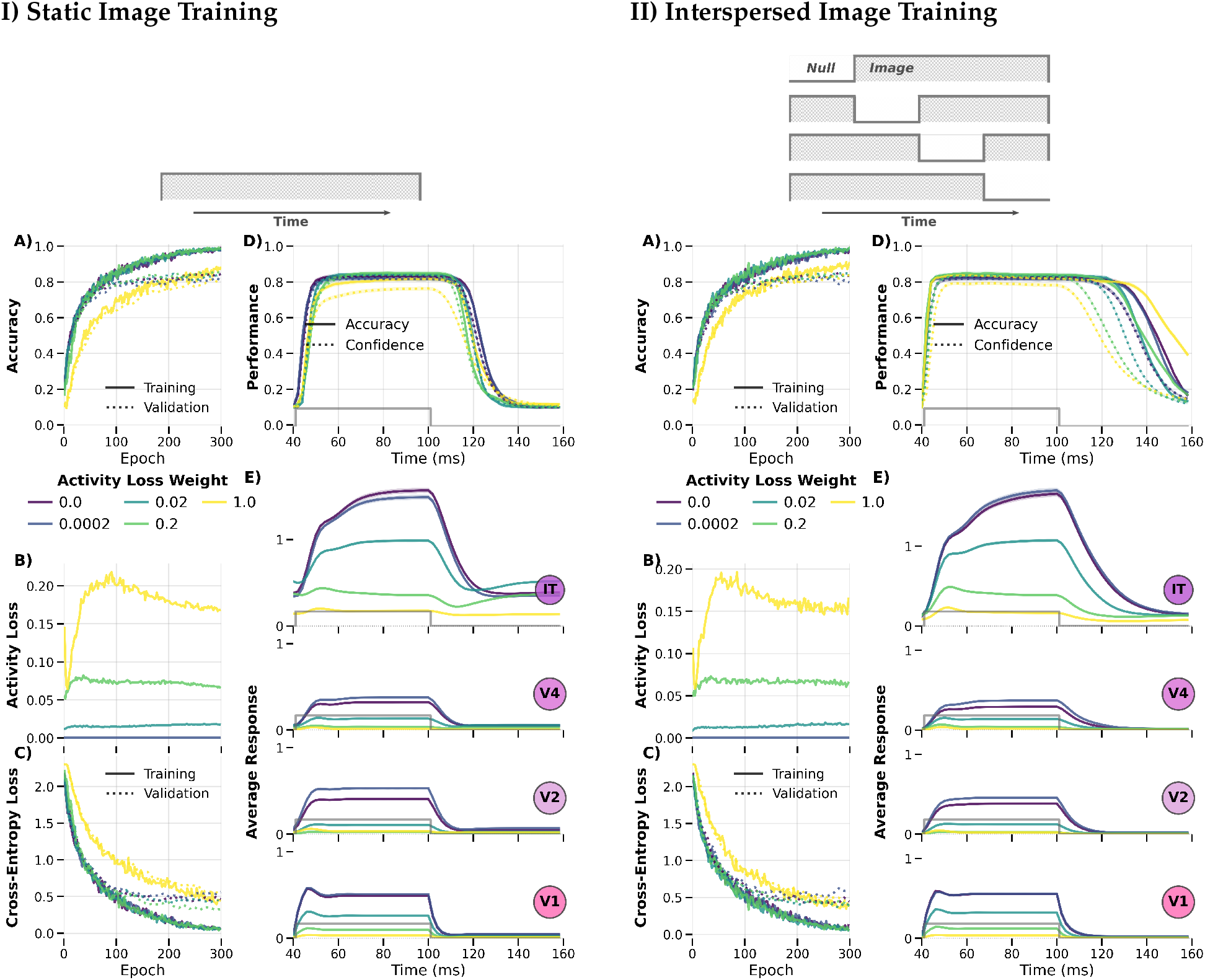
Training DyRCNNx8 models with different temporal presentation patterns and activity loss weights. **I)** Training on static images (pattern=`1`) extended for all timesteps (T= 40 ms). **II)** Training on images interspersed with null input (pattern=`1011` + shuffled) extended for all timesteps. **A)** Training (solid) and validation (dotted) accuracy across 500 epochs for different activity loss weights. **B)** Evolution of the activity loss component during training. **C)** Evolution of the cross-entropy loss component during training. **D)** Testing accuracy (solid) and confidence (dotted) on 40 ms idle null input (not shown) + 60 ms image input + 60 ms null input as indicated by the gray step function. **E)** Temporal dynamics of average activations for layers V1, V2, V4, IT for the same testing scenario as (D).

We observe that activity loss weights above a certain value ( ∼ 0.5) start to be detrimental to classification performance (panels A, D). Furthermore, small changes to the activity loss weight have visible effects on the model’s temporal response dynamics (panels E), by reducing the response magnitude and preventing response increases over the duration of a stimulus. As the application of an activity loss (L1 norm of activation tensors) encourages lower average activations, it also promotes stronger inhibitory recurrent weights (Supplementary Figures 23 and 24). The emergence of organized dynamics from metabolic constraints is consistent with findings that energy-efficiency objectives can cause RNNs to spontaneously develop predictive coding circuitry [50], paralleling how our activity loss promotes inhibitory recurrent organization without explicit instruction.

Besides training on static images, i.e. pattern `1` (Figure 6I), RCNNs can benefit from being trained on a combination of image and null inputs [51]. Therefore, we explore training on image presentations interspersed with a null input for a quarter chunk of the timesteps, the position of the chunk shuffled between batches, i.e. patterns `0111`, `1011`, `1101`, `1110`, (Figure 6II). Models trained on interspersed image presentation patterns show lower null responses and generally improved accuracy when using high activity loss weights.

These results establish that moderate metabolic constraints and static-image training provide stable dynamics without sacrificing accuracy. Models trained with interspersed null input produce cleaner null-period responses but require the chunk positions to be shuffled across batches to avoid learning the exact temporal sequence. We adopt static-image training with activity loss weight = 0.0002 as the default protocol for subsequent experiments unless otherwise specified.

### 3.4 Timestep Protocol: Duration, Loss Window, and Idle Period

#### Training Timesteps

During training the model is presented with input images for multiple timesteps. The number of timesteps used influences the stability of the learned temporal dynamics as well as training speed and memory allocation. Additional parameters such as the loss reaction time (see below) will also influence the total number of timesteps needed. In Figure 7A, we find that even a relatively small number of training timesteps (8 timesteps with 6 contributing to loss) are already sufficient to learn temporal dynamics and task performance that are stable for much longer testing time periods (60+ timesteps). Still, the choice of training timesteps needs consider the delays of skip, feedback, and recurrent connections, to allow signals from each connection type to reach the classifier (c.f. Figure 2).

**Figure 7:**
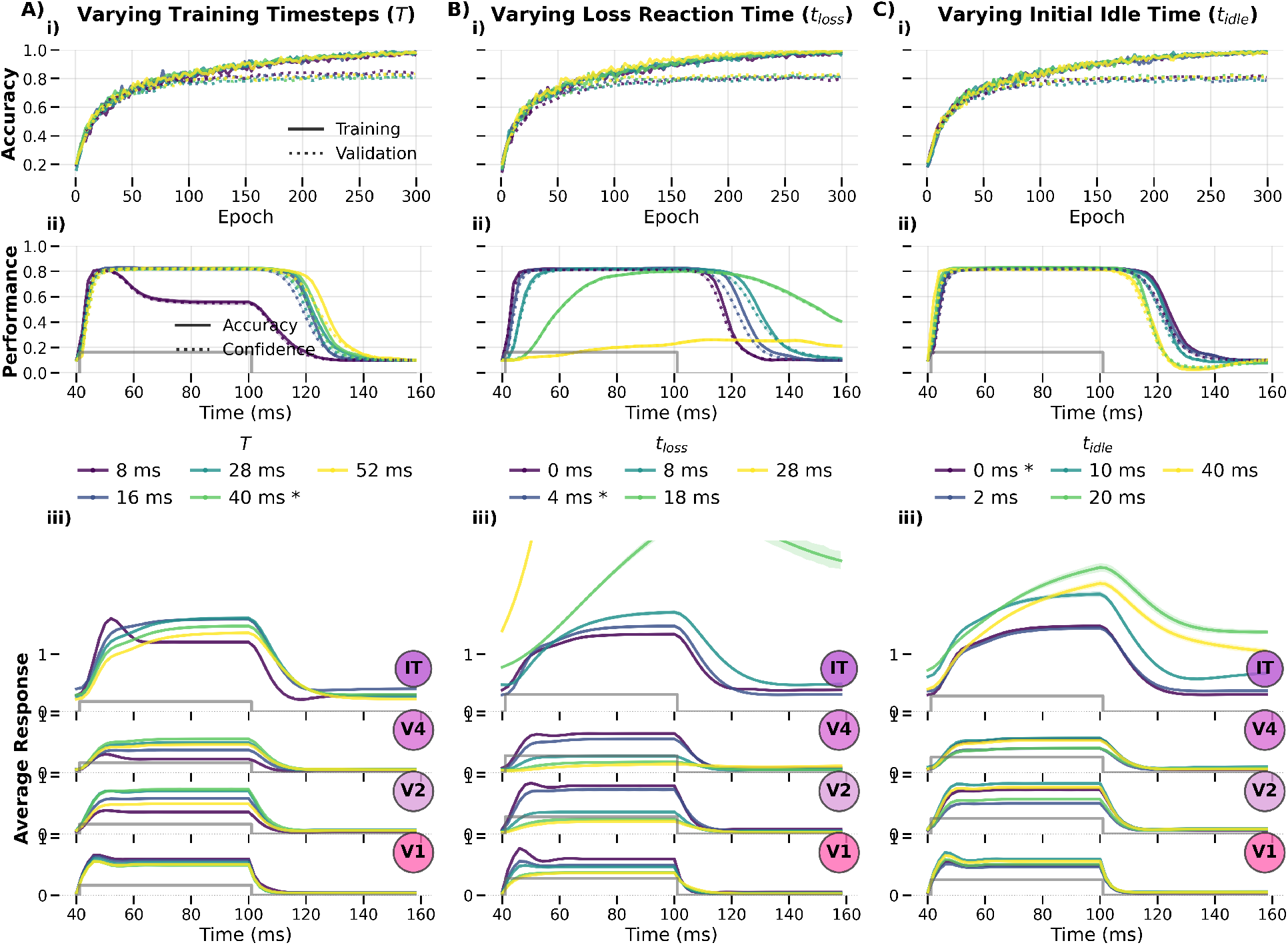
Effect of varying timestep parameters on learned temporal dynamics. **i)** Training and validation accuracy during training; **ii)** accuracy and confidence over simulation time during testing, the grey step function indicates the transition of stimulus presentation (0-60 ms) to null input (60-120 ms); Note that testing also includes an initial 40 ms idle period that is absent during training and can influence performance, e.g. for large *t*_*Loss*_ values in B)ii. **iii)** averaged response of each layer over time during testing, corresponding to (ii). Testing results in (ii) and (iii) are averaged over samples of each data category. Stars in the legend labels indicate the default parameter value. Column **A)** varies the training simulation time per input (default=40 ms); **B)** the initial period after stimulus presentation to ignore in the loss calculation (default=4 ms); **C)** the idle simulation time period with null input prior to the stimulus presentation (default=0 ms). Each parameter is shown in milliseconds assuming dt = 2 ms timesteps.

#### Loss Reaction Time

The evaluation of the training loss can be restricted to a subset of timesteps. In the literature, the exact number and placement of these timesteps varies across studies, including models that use all timesteps [10], the last timestep [15], or a number of timesteps in between [19]. We show in Figure 7B that the temporal window used for computing the training loss indeed strongly shapes learned dynamics, particularly at later layers. Using the entire output period (lossrt= 0 ms) produces rapid onset ramps that may overshoot before settling; restricting the loss to only the final few timesteps (lossrt ≥ 18 ms) leads to unstable runaway dynamics. Computing the loss over most timesteps promotes both stability and rapid classification. As noted by Spoerer et al. [10], this is biologically motivated, since object recognition is a continuously ongoing process. We find a small reaction-time offset (4 ms) prevents the optimization from over-prioritizing ramp-up speed at the expense of later stability.

#### Initial Idle Period during Training

We tested whether prepending idle timesteps with null input, as done in CordsNet [19], benefits our DyRCNN architecture. For the idle timesteps we choose zero tensors as the null input representing a neutral contrast-free input (this corresponds to the average pixel value, since the input images are normalized). As shown in Figure 7C, there is no evident benefit for training our model with initial idle timesteps (and may even cause instabilities). This is likely because the activity loss regularization provides sufficient constraint on activity levels from the start of training, eliminating the need for an idle stabilization period. This simplifies the training protocol while maintaining stable dynamics.

Together, these sweeps establish that stable dynamics emerge with as few as 8 training timesteps, that using most timesteps in the category loss promotes stability and prevents runaway dynamics, and that initial idle pretraining periods are unnecessary when activity loss is present. We adopt 20 timesteps (40 ms), loss reaction time of 4 ms, and no idle period as defaults.

### 3.5 Recurrence Architecture: Type, Integration Target, and Seed Consistency

#### Type of Recurrent Connections

We apply different types of recurrent connection strategies, realized by different types of convolutions as detailed in Section 2.3.1. All types of recurrence train to similarly high maximum validation accuracies: depthpointwise, localdepthwise (0.85) > full, pointdepthwise, none (0.82) > local, self (0.81). Figure 8A shows that they result in similarly shaped average layer response curves, with only the full recurrence model showing some deviations in form of an overshoot response in early layers upon stimulus onset. See the corresponding weight distributions in Supplementary Figure 20.

**Figure 8:**
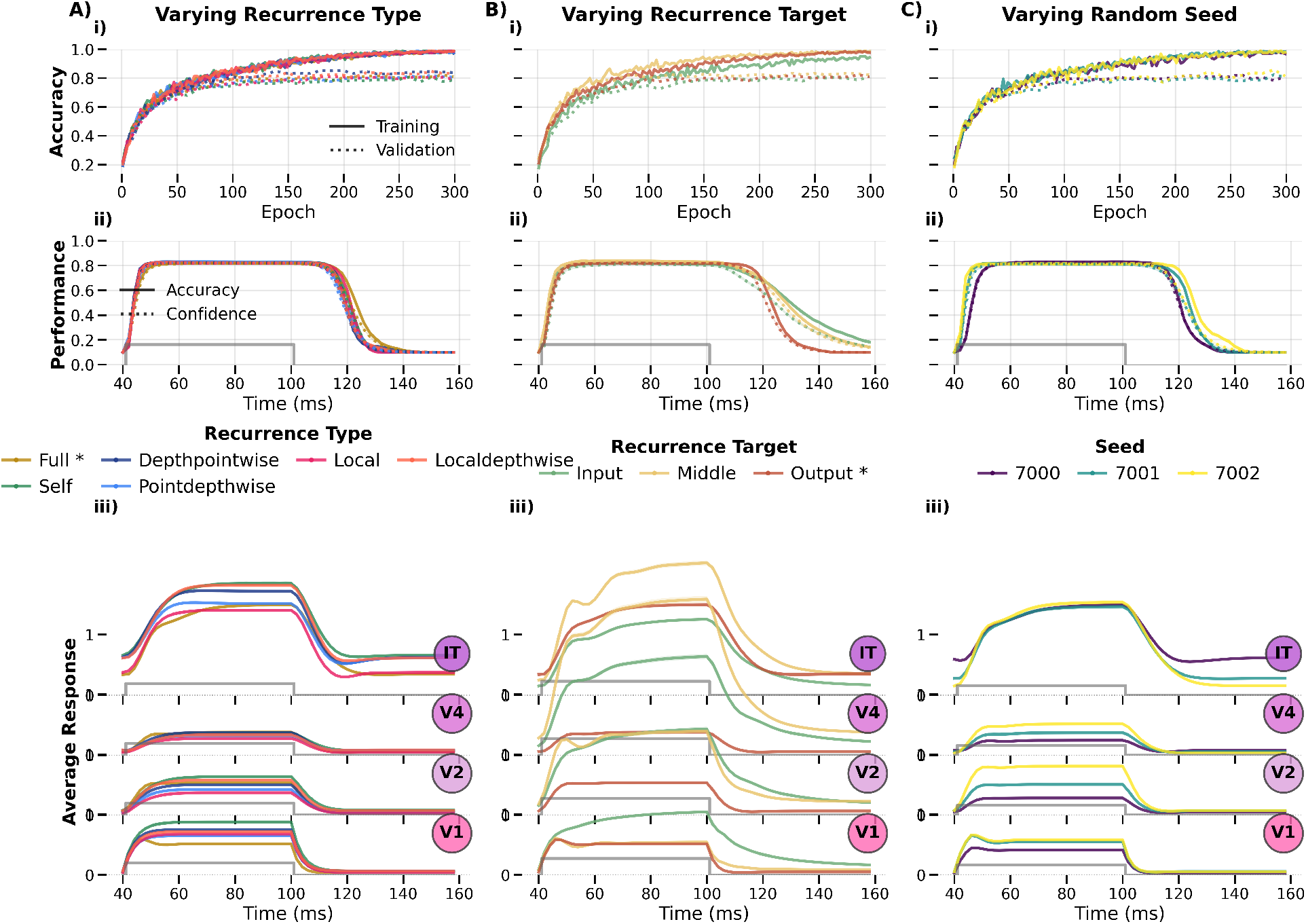
Impact of recurrence type and target on temporal dynamics, and variation across seeds. **i)** Training and validation accuracy during training; **ii)** accuracy and confidence over simulation time during testing, the grey step function indicates the transition of stimulus presentation (0-60 ms) to null input (60-120 ms); **iii)** averaged response of each layer over time during testing, corresponding to (ii). Testing results in (ii) and (iii) are averaged over samples of each data category. Stars in the legend labels indicate the default parameter value. Column **A)** varies the convolution-based recurrent connection strategy (default=full); **B)** the target where the recurrent signal is integrated relative to the two feedforward convolutions (default=output); **C)** the variation across random seeds for the default model.

#### Target of Recurrent Connections

Where the recurrent signal is integrated relative to feedforward computations (before convolutions: `input`, between them: `middle`, or after them: `output` see Section 2.3.3 for definitions) has a larger effect on temporal dynamics than the recurrence type itself (Figure 8B). The middle-target configuration produces the most qualitatively distinct dynamics, with pronounced onset transients and oscillatory settling behavior not seen in the input or output variants. See the corresponding weight distributions in Supplementary Figure 21.

#### Variability across Seeds

While other plots show the average over three random seeds, Figure 8C shows the temporal dynamics for the default model separately across different random initialization seeds. The qualitative shape of the response (onset transient, sustained plateau, and offset decay) is consistent across seeds, confirming that the dynamical patterns reported in other sections reflect architectural choices rather than initialization artifacts. Seeds differ primarily in response amplitude (layers V1-V4) and in the baseline activity level of the IT layer during null-input periods. Interestingly, IT baseline activity appears to correlate with classification dynamics near stimulus transitions: seeds with higher IT baseline activity show delayed accuracy onset and earlier accuracy decline at stimulus offset, hinting that a larger contrast between stimulus-driven and baseline activity in the readout layer supports sharper temporal classification boundaries. This observation warrants further investigation, as it may point to the importance of sparse baseline coding in the decision layer for temporally precise readout.

Taken together, these results establish that within the recurrence architecture, the integration target dominates over connectivity pattern in shaping temporal dynamics, and that the qualitative effects reported throughout this work are robust to random initialization.

### 3.6 Long-Range Connectivity and Input Protocols

#### Temporal Presentation Patterns

Extending the presentation-pattern analysis from Section 3.3 with patterns `1` and `1011`, Figure 9A also shows models trained with two and three quarter chunks receiving null input (pattern=`1001` and pattern=`1000`). We find that using presentation patterns that include null input improve the model’s null responses, both by reducing the magnitude of the layer responses and by maintaining the correct classification output for longer after offset of an image input. However, the pattern=`1000` is an exception because the model learns that any image input is always only one quarter of the timesteps long, and therefore shows response and performance artifacts when the testing stimulus is presented for longer. As before, chunk positions are shuffled between batches to prevent the model from learning the exact temporal sequence.

**Figure 9:**
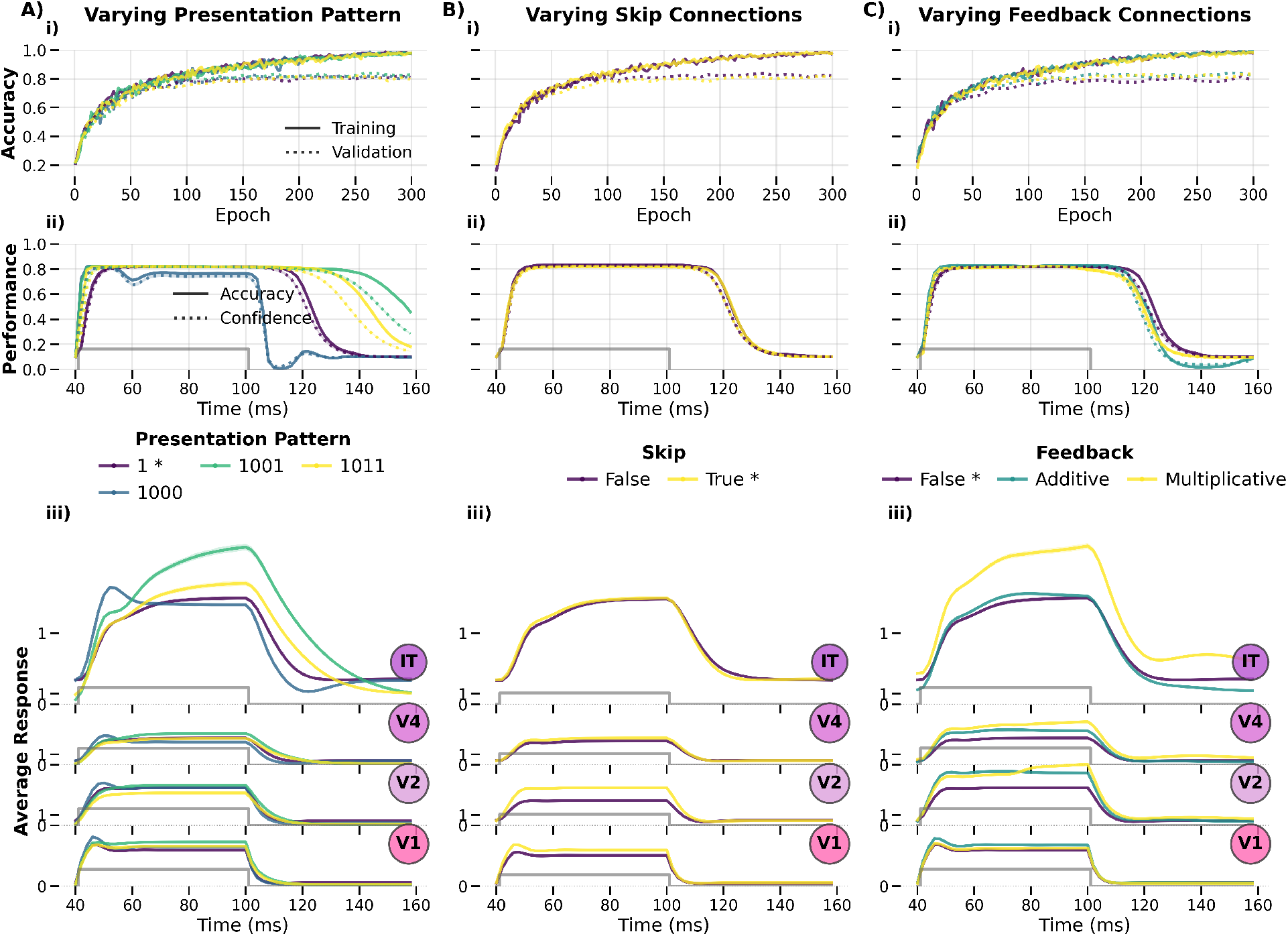
Impact of skip and feedback connections, and the temporal presentation pattern of input images during training. **i)** Training and validation accuracy during training; **ii)** accuracy and confidence over simulation time during testing, the grey step function indicates the transition of stimulus presentation (0-60 ms) to null input (60-120 ms); **iii)** averaged response of each layer over time during testing, corresponding to (ii). Testing results in (ii) and (iii) are averaged over samples of each data category. Stars in the legend labels indicate the default parameter value. Column **A)** varies the presentation pattern defining how many timesteps get null input instead of image input, `1`:static, `1011`:1/4 null, `1001`:1/2 null, `1000`:3/4 null, where the temporal sequence of null and image chunks are shuffled between batches (default=1); **B)** whether additive skip connection are present between V1->V4 and V2->IT (default=True); **C)** whether feedback connections between IT->V2, V4->V1 are present, and if their integration is additive or multiplicative to the feedforward signal (default=False).

#### Skip Connections

Adding skip connections to models, in particular deep models, is known to support efficient training by providing shorter error backpropagation paths along the computational graph to weights in earlier layers [45]. In our relatively shallow 8-layer DyRCNNx8 model, we observe little difference between models with and without skip connection in their training progress and their resulting temporal dynamics (Figure 9B). Skip weights remained small during training (final *σ* = 0.002 − 0.006), consistent with their order-of-magnitude smaller initialization (Supplementary Figure 22).

#### Feedback Connections

Feedback connections show minimal impact on model activity and classification performance, with the exception that multiplicative feedback modestly increases average IT activity (Figure 8C). This is plausible as the static image classification task provides little incentive for strong top-down signals, and feedback weights receive fewer gradient signals updates since they only contribute to the final few model outputs. Accordingly, feedback weights (initialization with 0.0 ± 0.001) tend to remain small after training, with additive feedback weights shifting slightly to the negative ( −0.00005 ± 0.0009), and multiplicative feedback weights shifting to positive values (0.0004 ± 0.0023) with a slightly broader distribution (Supplementary Figure 22).

We adopt static images with skip connections enabled and feedback disabled as defaults.

### 3.7 Temporal Parameters: Timescale and Conduction Delays

#### Time Constant *τ*

The time constant of the dynamical system (Figure 1) governs how quickly layer activity tracks its driven state. Figure 10A shows that larger *τ* values produce slower rise times in response to stimulus onset and slower decay following stimulus offset. Transient overshoots at stimulus onset, which are also commonly observed in neural responses, appear only for small *τ* values (< 9 ms), where the system responds fast enough to overshoot before settling.

**Figure 10:**
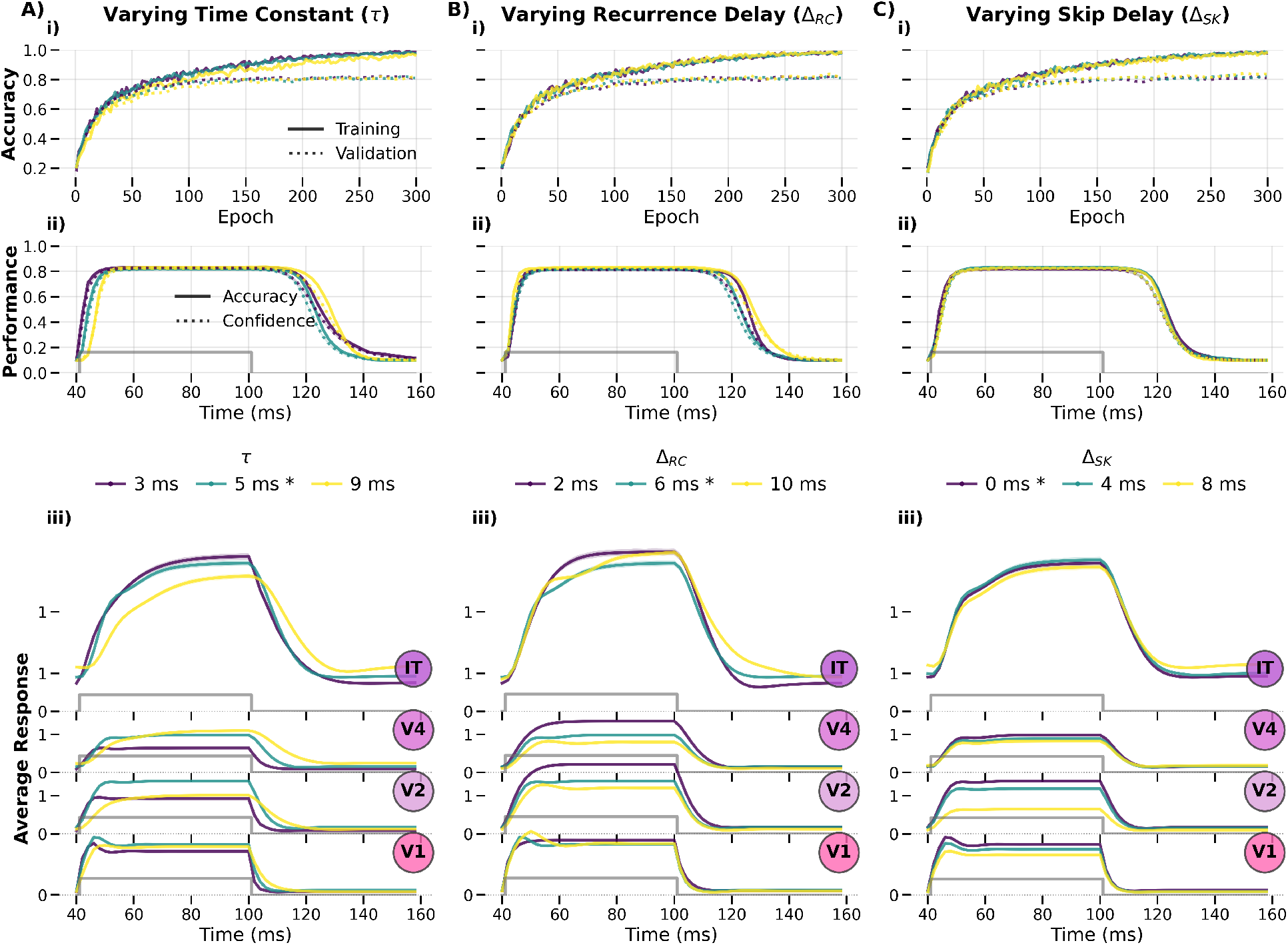
Effects of temporal parameters on network dynamics. **i)** Training and validation accuracy during training; **ii)** accuracy and confidence over simulation time during testing, the grey step function indicates the transition of stimulus presentation (0-60 ms) to null input (60-120 ms); **iii)** averaged response of each layer over time during testing, corresponding to (ii). Testing results in (ii) and (iii) are averaged over samples of each data category. Stars in the legend labels indicate the default parameter value. Column **A)** varies the time constant *τ* in the dynamical system equation (Figure 1) (default=5 ms); **B)** the temporal delay of recurrent connections (default=6 ms); **C)** the temporal delay of skip connections (default=0 ms in engineering time, is equivalent to Δ_*FF*_ in biological time).

#### Recurrence Delay Δ_*RC*_

The recurrence delay determines how far back in time the lateral recurrent signal originates. Figure 10B shows that larger Δ_*RC*_ values produce more pronounced adaptive behavior, including slight oscillatory transients resembling the overshoots seen with small *τ*. This is expected since a longer delay in the recurrent feedback loop increases the lag between excitation and its self-regulation. Both *τ* and Δ_*RC*_ values have minimal effect on classification performance in the tested range.

#### Skip Connection Delay Δ_*SK*_

Figure 10C shows that the skip connection delay has little to no effect on temporal dynamics or classification performance across the tested range. This is consistent with our earlier observation that skip connections carry smaller weights in this architecture (Supplementary Figure 22), limiting their influence regardless of delay.

Classification accuracy is largely insensitive to these parameters in the tested ranges, indicating robustness to timescale variation. We adopt *τ* = 5 ms, Δ_*RC*_ = 6 ms, and Δ_*SK*_ = 0 ms as defaults.

### 3.8 Summary of Parameter Selection

In the previous sections, we showed a range of parameters under which models can produce good task performance and reasonable activity profiles, while also considering the computational cost of different model settings. Through this systematic exploration of the modeling parameter space, we established the configuration used for all DyRCNNx8 models in the biological validation experiments: 20 training timesteps with dt=2 ms, no initial idle period, skip but no feedback connections, and the temporal parameters specified in Table 1. These choices balance biological plausibility with computational efficiency while producing stable, interpretable dynamics.

### 3.9 Computational Performance Benchmarking

Beyond the 52% speedup demonstrated with CORnet-RT, we provide computational performance analysis showing how different modeling choices affect training speed, GPU memory usage, and model complexity (Table 2).

**Table 2:**
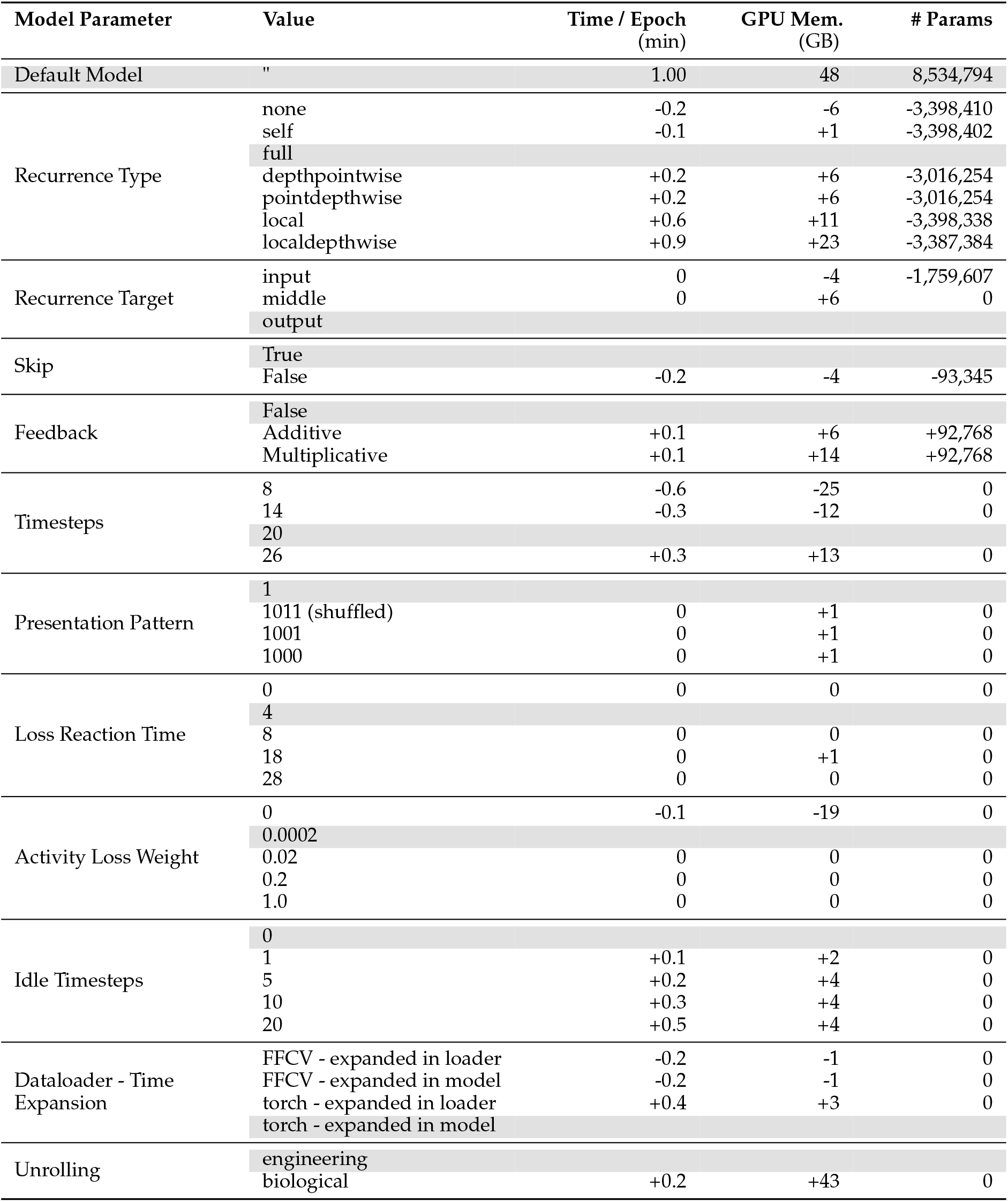
Computational resource demands for different modeling choices. Values show the change relative to the default model configuration (`DyRCNNx8`, recurrence type `full`, target `output`, 20 timesteps with 2 ms resolution, pattern `1`, skip connections enabled, no feedback, loss reaction time 4 ms, activity loss weight 0.0002, idle timesteps 0) trained in ImageNette. GPU memory reported as the 90^th^ percentile of `system.gpu.0.memoryAllocatedBytes` (PyTorch-allocated memory during training; robust to warm-up and validation-phase drops). Time per epoch computed from the slope of `runtime` vs. `epoch`. All runs use FP32 precision, Lightning 1.9.5, NVIDIA A100-SXM4-80 GB, batch size 192, averaged over 3 seeds. Gray rows denote the default value for that parameter category. Within-group standard deviation ≤ 0.5 GB for all rows.

#### Architectural choices with counterintuitive resource demands

Several architectural choices produce resource demands that are not obvious from parameter counts alone. Targeting recurrent connections at the middle stage of two-stage convolutions (`rctarget=middle`) increases GPU memory by ∼ 6 GB relative to the output target despite identical channel dimensions; profiling reveals that the autograd engine must retain both the first convolution’s output and the recurrence output for the second convolution’s backward pass, whereas output-target recurrence runs last and avoids this double-storage. Biological unrolling (`tff`=10 ms, `tsk`=20 ms) nearly doubles GPU memory relative to the engineering default, as the extended delay buffer stores ∼ 22 ms of hidden-state activations across all four layers. In contrast, replacing the the convolution in full-recurrence with the piecewise scalar multiplication of self-recurrence shows no meaningful resource benefit. Disabling the activity loss (`activityloss`=0) reduces GPU memory by ∼ 19 GB, as no activity norms need to be tracked during training. Training time and memory scale predictably with the number of timesteps, skip connections, loss reaction time, and idle timesteps.

#### Dataloader comparison

The GPU-optimized FFCV (Fast Forward Computer Vision) dataloader can achieve 1.2–7× epoch-speed improvements over the default PyTorch dataloader by pre-processing and storing data in a format optimized for fast loading with minimal CPU overhead [27]. However, FFCV uses its own data compression and augmentation pipeline, so training results can diverge from the PyTorch baseline. For our setup, FFCV provided only a ∼ 16 % speedup and a ∼ 4 % GPU-memory reduction (Table 2). The gain is modest because the 20-timestep forward pass makes the workload compute-bound rather than I/O-bound; additionally, the ImageNette dataset (∼ 9,500 images, 10 classes) is small enough for the default PyTorch dataloader with 4 workers to handle it efficiently. FFCV is expected to provide larger benefits for ImageNet-scale training or multi-GPU distributed setups. Beyond throughput, we found that FFCV-trained models required more epochs to reach comparable accuracy but were less prone to overfitting. When comparing temporal dynamics of FFCV- and PyTorch-trained models at matched accuracy, notable differences in learned representations can persist, likely reflecting the different data-augmentation implementations used by the two backends. We therefore recommend evaluating dataloader choices carefully in light of the specific research question and resource constraints.

### 3.10 Evaluation of Biological Plausibility

Having explored the various design impacts on training and activity, we now evaluate the extent to which the resulting models exhibit biologically realistic dynamics that match both neural data and behavioral observations. In particular, we focus on different recurrence types and targets as well as different training presentation patterns and activity loss weights.

#### 3.10.1 Stability and Response to Null Input

Biological neural networks maintain stable baseline activity and respond appropriately to the absence of input. To verify these properties in our models, we examined network dynamics both during prolonged input presentation beyond the training timesteps and following stimulus offset.

Figure 11 demonstrates that trained models maintain stable activity levels far beyond the range of training timesteps without exhibiting runaway excitation or complete silence, both of which would indicate biologically implausible dynamics. Furthermore, network activity returns to stable baseline levels following stimulus offset, matching the behavior observed in cortical recordings. These properties emerge in our models naturally without the model ever being exposed to null input, without explicit normalization, and without the need for an activity loss penalty (as shown by the response of activity loss weight = 0).

**Figure 11:**
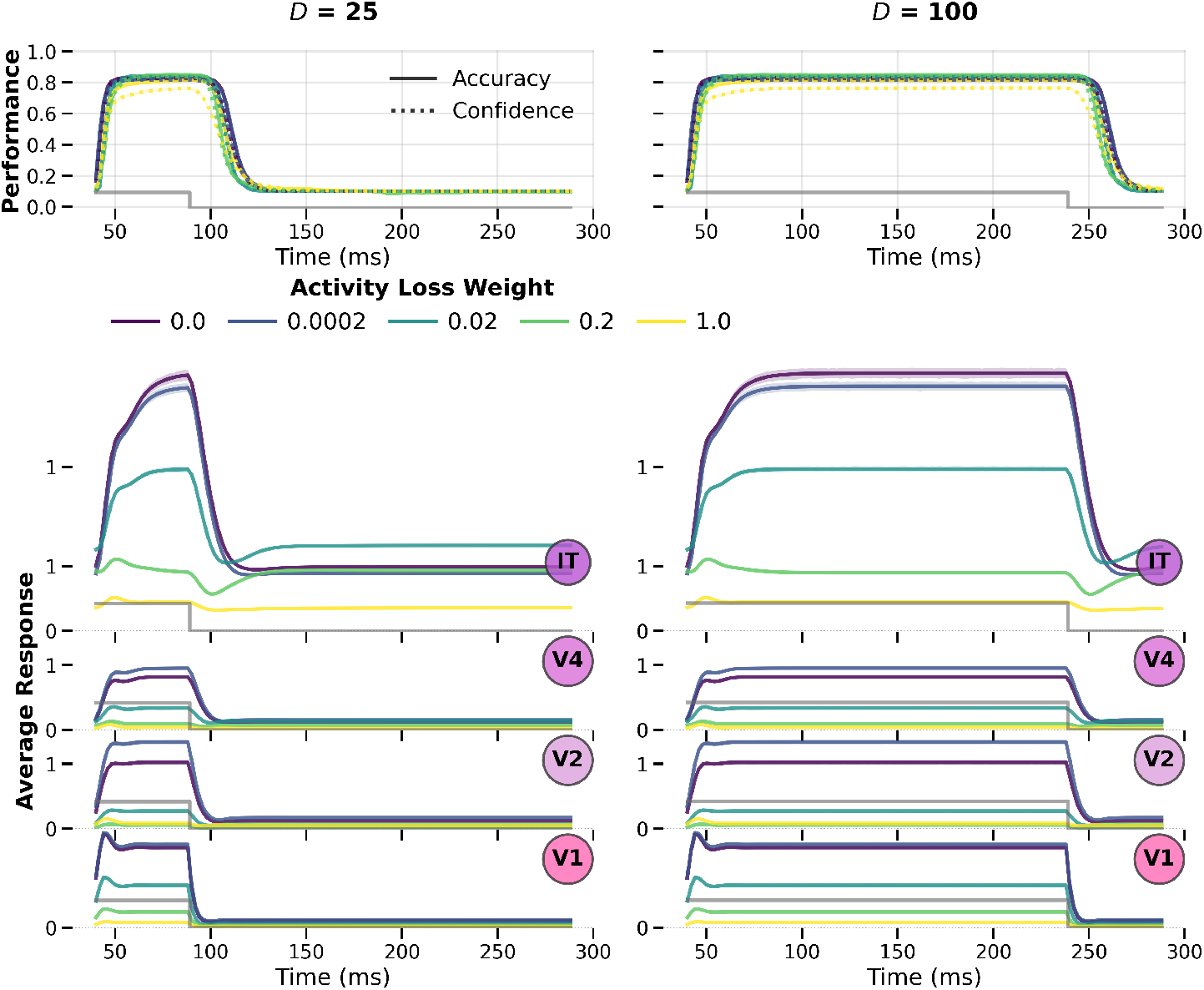
Models exhibit stable dynamics and appropriate null responses. Our DyRCNNx8 model with default parameters (see Table 1) trained on 20 timesteps of static images is tested on 125 timesteps preceded by an additional 20 timesteps of idle null input. The left column **I** shows the model’s response to *D* = 25 timesteps of image input followed by null input for the remaining inputs, and the right column **II** shows the response to *D* = 100 timesteps of image input. **A)** shows the test accuracy (*solid*) and confidence (*dotted*) over time. Different hues indicate model versions that were trained with different weighting factors for the activity loss. The timesteps of image presentations are indicated by a grey step function. **B)** shows the corresponding average layer-wise responses. Supplementary Figure 25 shows the same results for training with interspersed images, and Supplementary Figure 26 for different recurrence targets.

#### 3.10.2 Comparison to Neural Data

In support of our goal of building and evaluating models based on their fidelity to biological neural dynamics, we analyze model dynamics using methods inspired by neural data analysis. Following Groen et al. [52], we test whether our models exhibit a well-known function of visual recurrence: temporally delayed normalization. This includes adaptation to repeated stimuli, sublinear summation of responses to stimuli of increasing duration, and contrast-dependent reaction time. Groen et al. [52] compute neural dynamics from broadband power (50-200 Hz) averaged across multiple ECoG electrodes per visual area, trials, and patient. We emulate an analog proxy measure of neural activity by averaging the model’s unit activations per layer and timestep (across stimuli presentation including correct and incorrect classifications). Using this analogy, in Figure 12 we explore how the type of recurrence impacts temporal neural responses. For these analyses, we employed the interspersed image presentation pattern `1011` during training as it was found to improve null response behavior Section 3.3. It should be noted that while the ECoG recordings show activity in visual area V1, Groen et al. [52] report qualitatively similar response characteristics for areas V2 and V3. In our architecture, V1 receives direct retinal input while V2 represents the first recurrent layer, and both map plausibly to the early visual cortex. Therefore, we choose to show the model’s V2 layer responses for the interval and duration experiments and the model’s V1 layer responses for the contrast experiment, which show a better fit to the data. We also present the corresponding results for the model’s V1/V2 layer responses in Supplementary Figure 15.

**Figure 12:**
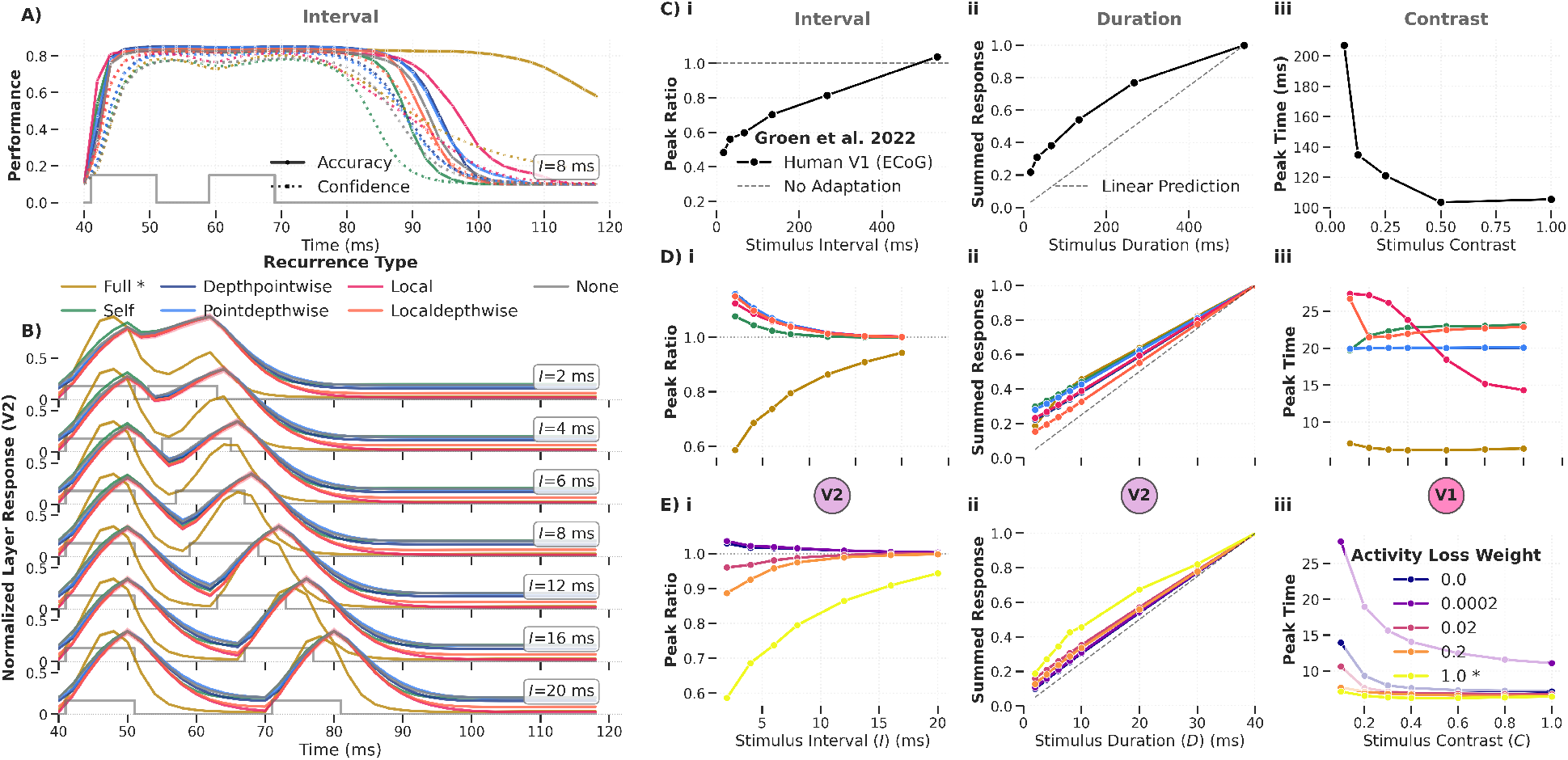
Temporal RCNN dynamics for different recurrence types in response to three test scenarios: varied inter-stimulus interval (i), stimulus duration (ii), and contrast (iii). The testing input is a static image shown for a selection of timesteps alternated with null input (zero tensor) which is indicated by the gray step-function in the subplots. The model (DyRCNNx8) was trained on ImageNette for 20 timesteps and is tested on 60 timesteps. **A)** Accuracy (solid) and Confidence (dashed) performance of the model variants over the testing time for repeated 10 ms image input separated by a *I* = 8 ms interval. **B)** Average responses of layer V2 to repeated image input with varied intervals *I*. **C** shows experimental data from ECoG recordings in human V1 [52] illustrating actual neural characteristics, and panels **D)** and **E)** the corresponding characteristic seen in our model variants with different recurrent types (default: `full`) and different activity loss weights (default: 1.0). The panels **i** show the measured height ratio of the two peaks in response to the repeated stimulus with respect to the inter-stimulus interval values (*I*), illustrating an adaptation effect. Analogously, panels **ii** show the activity summed over time in response to stimuli of different duration (*D*), illustrating sublinear summation. Panels **iii** show the time-to-peak in response to a stimulus with varying contrast, illustrating delayed reaction times to weak stimuli.

##### Response to repeated stimuli with varying interval

Paralleling the experimental setup of [52], Figure 12A shows activity in response to a 10 ms stimulus that is repeated after an 8 ms interval of null input. Figure 12B shows the corresponding average layer V2 responses for intervals increasing from 2 ms to 20 ms. The experimental recordings from Groen et al. in Figure 12Ci show that for stimuli repeated in quick succession the response peak for the second stimulus is reduced, i.e. the peak ratio is < 1, but the second peak recovers to the same height as the interval increases. As shown in Figure 12Di, this adaptation effect emerges in our model only for the `Full` recurrence type, and as shown in Figure 12Ei for large activity loss weights. Other types of recurrence and small activity loss weights rather cause the second response peak to be larger than the initial response peak. This shows how replicating biological dynamics requires careful parameter choices.

##### Response to stimuli with varying duration

Presenting a stimulus typically triggers a strong response around the stimulus onset that decays to a lower plateau activity for the remainder of the stimulus duration. Therefore, measuring the summed activity in response to stimuli of increasing duration shows a sublinear summation effect, as demonstrated experimentally by Groen et al. and displayed in Figure 12Cii. In our model, this effect is much smaller but again most pronounced for models with full recurrence and a larger activity loss weight, as shown in Figure 12Di-Ei. Similarly, [53] find that only CNNs with lateral recurrence, not feedforward models, capture human-like performance across stimulus durations.

##### Response to stimuli with varying contrast

Lastly, Groen et al. also demonstrate a decrease in reaction time, i.e., time to peak, with increasing image contrast (Figure 12Ciii). Different than for the interval and duration effects, our models show a contrast-dependent reaction time only for either local recurrence or for full recurrence with minimal or no activity loss (Figure 12Diii-Eiii).

Groen et al. model the neural characteristics of adaptation, sublinear summation, and contrast-dependent reaction time with a delayed normalization (DN) model (c.f. panel C in their Figures 7, 8, 9 respectively). Notably, our model architecture does not include any normalization operation. Instead, any normalization effect relies on the recurrent connections. This demonstrates that continuous-time recurrent dynamics can naturally give rise to key temporal phenomena observed in cortex without requiring explicit divisive normalization operations. Additional temporal dynamics figures for the complementary V1-V2 results as well as for different recurrence targets are shown in the Supplementary Figures 15, 16, and 17.

#### 3.10.3 Robustness to Noise

A hallmark of biological vision systems is their robustness to noisy or degraded input. Recurrent connections have been proposed as a mechanism that could enhance robustness by enabling temporal integration and contextual modulation. To test whether our models exhibit this property, and to link model behavior to human psychophysics, we evaluated the performance of models trained on static images (without any noise) with different recurrence targets and activity loss weights on images corrupted with varying levels of Gaussian noise.

Figure 13 demonstrates that noise robustness depends strongly on where a recurrent signal is integrated within a layer. Integrating recurrence before (*input*) or between (*middle*) the feedforward convolutions significantly improves accuracy on noisy images compared to integrating it after the convolutions (*output*); the *output* variant offers only marginal improvement over an equivalent feedforward model trained without recurrence (*None*). Disabling recurrent connections at test time on models trained with recurrence reduces performance across all variants (△ markers in Figure 13A, dashed lines in panel B), confirming that the observed robustness gains depend on active temporal integration rather than on static architectural differences alone.

**Figure 13:**
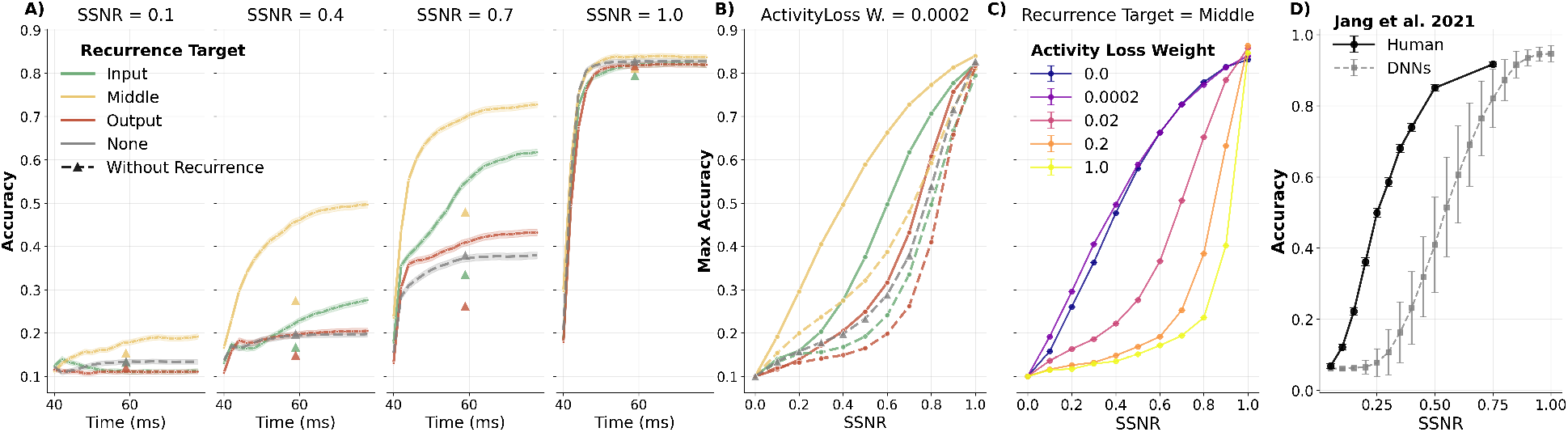
Some recurrence types show noise robustness more similar to human behavior. Model variations with different recurrence target locations (*colored*: input, middle, output), each trained separately on static images with activity loss weight = 0.0002, each with three different seeds, are tested on images corrupted with Gaussian noise with varying signal-to-signal-plus-noise ratios (SSNR). **A)** Categorization accuracy traces over time, averaged over image samples and seeds. Shaded areas indicate the standard error of the mean. △ *markers* indicates the maximum accuracy of the same models with disabled recurrent connections. **B)** Maximum performance (over time) as a function of noise intensity (SSNR). Error bars indicate standard deviation across seeds. The *dashed* lines indicate the maximum accuracy of models with disabled recurrence. **C** Same as panel B, but for middle-target variant trained with different activity loss weights. **D)** Human (*black, N* = 20) and DNN accuracy (*grey*) as a function of SSNR for a 16-class image classification task, replicated from [54] (c.f. their Figure 2a). DNN accuracy is the average of AlexNet, VGG-F, VGG-M, VGG-S, VGG-16, VGG-19, GoogLeNet, and ResNet-152.

Comparing to human data [see panel D with data from 54], Figure 13B shows that the *middle*-target variant best captures human performance trends. Figure 13C further reveals that higher activity loss weights progressively reduce noise robustness in our models. Supplementary Figure 19 shows that with low activity loss and middle-target the *full* recurrence still performs best. This presents an unexpected dissociation: the activity loss that promotes the biologically realistic temporal dynamics (Section 2.2) also reduces the model’s capacity to handle noisy input. This pattern, where recurrence alone does not automatically confer robustness but instead depends on its specific configuration and interaction with other network properties, aligns with recent work showing that top-down feedback improves noise robustness only when combined with stochastic regularization [55]. A parallel pattern appears in our own feedback results: feedback connections show no systematic noise-robustness benefit across seeds, with multiplicative feedback being only marginally preferable to additive (Supplementary Figure 18). In both cases, the functional impact of recurrence depends not on its mere presence but on the specific architectural and training context in which it operates.

### 3.11 Framework Generality: Reference Model Reimplementation

Having systematically explored the DyRCNNx8 parameter space and documented the biological plausibility of the resulting models, we now demonstrate the framework’s generality and flexibility. We show that DynVision can precisely recreate established models from the literature, validating the toolbox’s correctness while revealing implicit modeling choices that were previously hidden or underspecified in the original implementations. We validated the exact replication of model function by asserting that each layer’s response is numerically equivalent between the original and the reimplemented model when presented with the same input.

#### 3.11.1 CORnet-RT

CORnet-RT [46] represents a successful approach to biologically-inspired recurrent vision models, optimized for Brain-Score performance. We reimplemented CORnet-RT within DynVision and initialized it with its ImageNet-pretrained weights.

CORnet-RT has an architecture with two convolutions in each of its four layers (V1, V2, V4, IT), each followed by a GroupNorm and a nonlinearity. Its recurrence adds the layer output unchanged to the feedforward signal before the second convolution. This corresponds in our nomenclature to a self-recurrence with middle-target and recurrence delay of one timestep. In the original code, the way the forward function loops over the layers also introduces a feedforward delay of one timestep, meaning a signal needs four timesteps to reach the classifier. Only this timestep is then used for the classification loss. In total, the training uses only 5 timesteps. In the reimplementation, these modeling choices are made explicit as parameter configuration that can be modified without changing the model code. Notably, recent work has extended the CORnet family with fMRI-based training objectives to further improve brain alignment [56].

Column I of Figure 14 presents the test responses of the reimplementation on the 10-class ImageNette dataset. Panel A shows its performance peaking for exactly the 4th timestep to the accuracy reported in the original publication [ 60% in 15]. Panel B shows the corresponding layer response dynamics.

**Figure 14:**
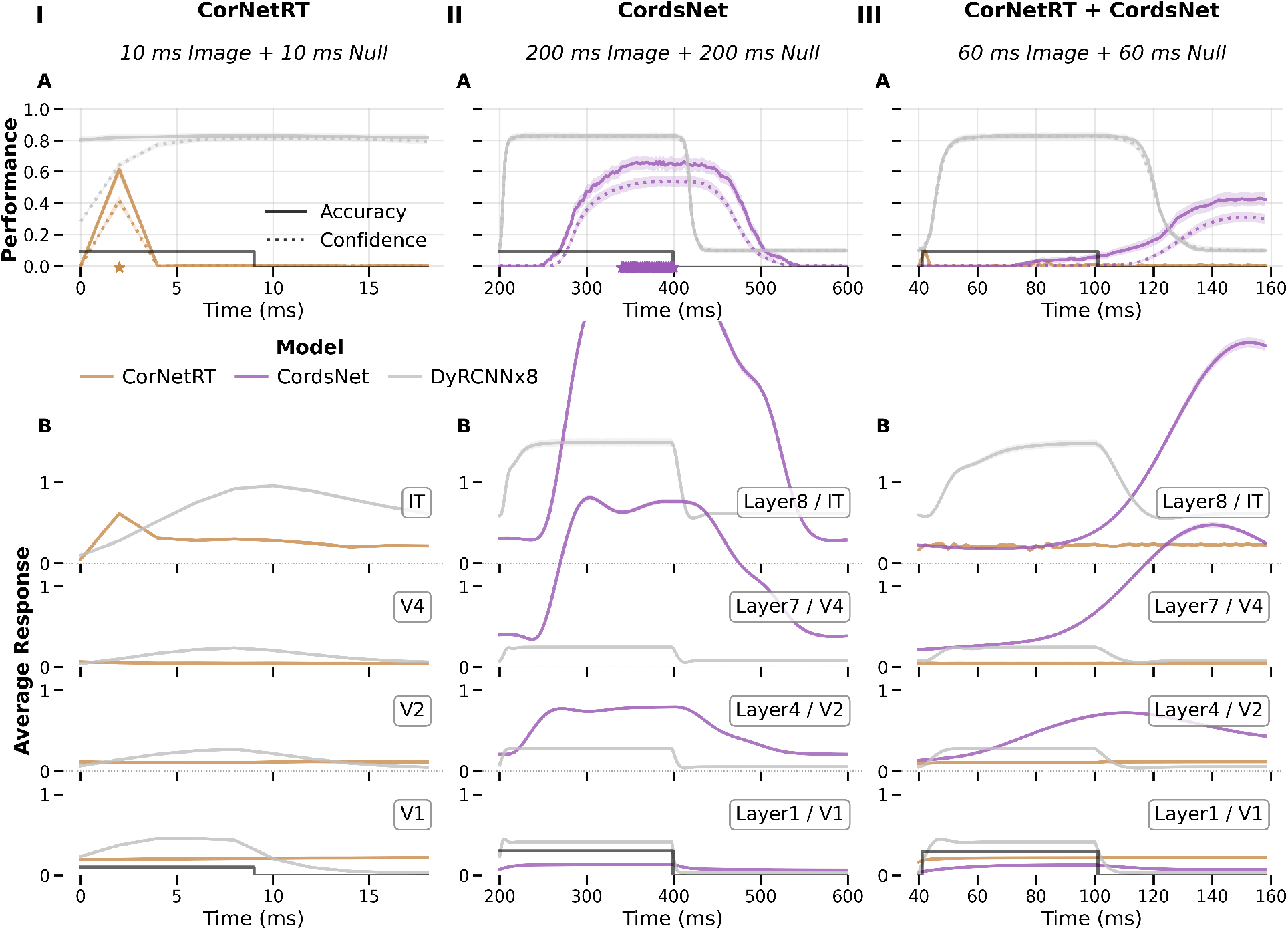
Equivalent reimplementation of CORnet-RT [15] and CordsNet [51] in DynVision. The CORnet-RT (*amber*) and CordsNet (*purple*) reimplementations are numerically equivalent to their original, initialized with ImageNet-pretrained weights and tested on ImageNette (10-class ImageNet subset) with a temporal presentation (indicated by *dark-grey* step function) representing their respective training setups (timestep *dt* = 2 ms). Corresponding DyRCNNx8 responses are shown in *light-grey* for reference. **I)** CORnet-RT (Δ_*FF*_ = 0 ms) tested on 10 ms image followed by 10 ms null input. **II)** CordsNet (Δ_*FF*_ = 2 ms, Δ_*SK*_ = 2 ms) run with 200 ms idle null input (not shown) followed by 200 ms image input followed by 200 ms null input. The layers 1, 4, 7, and 8 are picked as representatives from its 8-layer architecture. **III)** Both models tested on a non-native timescale of 40 ms idle null input (not shown) followed by 60 ms image input followed by 60 ms null input. **A)** Accuracy (*solid*) and confidence (*dotted*) over testing time. The *star* marker(s) indicate which timesteps were used for the category loss during training. **B)** Averaged layer responses over testing time.

Column II of Figure 14 shows the same model in response to a testing setup that is slightly different from how the model was originally trained in that it presents idle null inputs before the image inputs. We find that CORnet-RT loses its classification capabilities when the image presentation is prepended even with a single null input, demonstrating brittleness in its temporal processing.

To further evaluate the efficiency of the reimplementation, we also retrained both the original and reimplemented model versions on ImageNette (a 10-class ImageNet subset). Our reimplementation achieved identical classification accuracy while delivering a 52% speedup: the original code required 13.51s per epoch while the DynVision implementation required only 8.86s per epoch on an NVIDIA A100 GPU (averaged over the first 80 epochs). Notably, our implementation computes the loss for all 5 timesteps rather than only the last timestep, performs more extensive logging, and applies more comprehensive data augmentations, extending the functionality of the original code. This successful reproduction validates the toolbox’s correctness and foreshadows the computational efficiency gains detailed in the subsequent benchmarking section.

#### 3.11.2 CordsNet

CordsNet [19] implements continuous-time dynamics (*dt* = 2 ms) with an approach philosophically similar to DynVision, making it an important reference point. Reimplementing CordsNet within our framework revealed several implicit design choices in the original implementation: The model implements a “full” recurrence that is integrated at the layer output in the next timestep, i.e., Δ_*RC*_ = 2 ms. Their model layers are processed in reversed order (latest to earliest) for each timestep, which effectively introduces a feedforward delay and skip delay of one timestep as any layer output is passed to the subsequent layer only in the next timestep iteration. During training, the model is presented with 100 initial idle timesteps, followed by 100 timesteps showing the image, and another 100 timesteps of null input. As the image is only shown for the first half of the (non-idle) timestep, this realizes a “`10`” presentation pattern. The first 70 (non-idle) timesteps are ignored for the classification loss, setting the loss reaction time to 140 ms. Note that the model’s connection delays Δ_*FF*_ = 2 ms and Δ_*SK*_ = 2 ms are presumably not designed to represent “biological time” as it is not biologically plausible for skip connections to have the same delay as single-layer feedforward connections [57, 58]. This setup cannot be equivalently transformed to true engineering time with Δ_*FF*_ = 0 ms either, because with the transformation explained in Section 2.2.3 the skip delay would become negative Δ_*SK*_ = −2 ms.

Column II of Figure 14 shows the model’s test responses using a temporal presentation that is equivalent to its training setup. Panel A shows the corresponding accuracy time curve, also peaking around 60% as reported in the original publication. Panel B shows the response dynamics of four representative layers of its eight total, with increasing response amplitude in later layers. Notably, the model doesn’t use an activity loss to penalize high activations but does have a loss to incentivize the return to baseline for null inputs.

Column III of Figure 14 presents the model again with a temporal testing setup that varies from its training setup. Its significantly longer rise time in response to stimulus onset is indeed more similar to neural response time in corresponding cortical areas. However, this is not a result of feedforward delays, which are minimal with Δ_*FF*_ = 2 ms per layer, but most likely stem from a longer dynamical systems timescale *τ* = 10 ms and the omission of the first 70 timesteps for the category loss. These parameter choices increase the computational costs associated with training and running the model.

It needs to be highlighted that the amplitude of the performance curves of our DyRCNNx8 model in Figure 14A (in grey) cannot be directly compared to the CORnet-RT and CordsNet model, which were trained on the 1000-class ImageNet dataset. This is because, to facilitate model exploration, our models were only trained on the ImageNette 10-class subset that is used for testing here.

The successful reimplementation of models with different architectural philosophies from the literature demonstrates that DynVision provides a flexible substrate for exploring diverse biological RCNN architectures. The explicit parameterization system makes previously hidden modeling choices visible and systematically explorable, as demonstrated throughout our DyRCNN analyses.

## 4 Discussion

Biologically-realistic recurrent convolutional neural networks occupy a productive middle ground in computational neuroscience: abstract enough for task-level behavioral comparison, yet sufficiently detailed for meaningful comparison with neural recordings at the population level. However, the mixed results across RCNN studies, often attributable to underspecified parameter choices and differing evaluation protocols, have made it difficult to draw general conclusions about the nature of recurrence in visual processing. Here, we share with the research community DynVision, a toolbox to address this by making the implicit modeling decisions that vary across studies explicit, configurable, and systematically and efficiently explorable. Our exploration of architectural choices reveals how qualitative patterns in temporal dynamics are highly sensitive to choices that are rarely discussed: the target location of recurrent integration, the temporal window used for loss computation, and the relative weighting of metabolic constraints. These choices shift quantitative performance and determine which biological phenomena a model exhibits.

Our results demonstrate that recurrent connections can realize temporal dynamics that are typically attributed to explicit normalization mechanisms. Specifically, models with full lateral recurrence targeting the layer output, trained with the strongest metabolic constraints we explored (activity loss weight ≈ 1, at the upper limit before classification accuracy degrades), exhibit sublinear temporal summation and adaptation to repeated stimuli, which are hallmark properties of cortical responses typically modeled using a delayed divisive normalization mechanism [52, 59]. Notably, our architecture contains no explicit normalization operation. Instead, recurrent weights converge to slightly negative values on average (Supplementary Figure 20), suggesting that the network learns an effectively inhibitory lateral recurrence that stabilizes feedforward activity. This inhibitory organization emerges from a standard image classification objective, being trained on a static image presentation interspersed with a null input (presentation pattern = `1011`) with no task-specific incentive to produce normalization-like dynamics. This finding directly addresses a question raised earlier: biologically plausible temporal dynamics can emerge from simple recurrent connections combined with metabolic constraints, without requiring explicit normalization mechanisms. It also aligns with theoretical work showing that recurrence alone can implement divisive normalization [20, 60, 61], and with empirical evidence that temporally filtered normalization explains interactions in V1 that static normalization cannot capture [62]. We extend these principles to task-trained networks operating at millisecond temporal resolution. However, this model variant (recurrence target = output, activity loss weight = 1, presentation pattern = `1011`) cannot replicate the contrast-dependent reaction time and robustness to noisy input observed in human experiments.

Strikingly, we find a different architectural configuration with full recurrence targeting the middle of a layer’s computations, and trained with minimal metabolic constraints on purely static images, that produces a qualitatively distinct dynamic regime. This variant shows much weaker temporal normalization but achieves substantially improved robustness to Gaussian noise, approaching human-level performance curves [54]. It also better fits the decreased neural reaction time to low contrast inputs observed in cortical recordings [52]. The dissociation between these two regimes suggests that the activity loss that promotes biologically realistic temporal normalization appears to encourage sparse coding that reduces redundancy, which in turn diminishes robustness to noisy inputs. Conversely, the middle-target configuration, which integrates recurrent signals between feedforward convolutions, may support richer spatial integration that aids denoising. This trade-off suggests that recurrence may serve functionally distinct roles depending on architectural context: one regime favoring temporal regularization through normalization, another favoring spatial integration for noise tolerance. More broadly, understanding whether the two functional regimes we identify map onto distinct cortical circuit motifs, for instance, laminar feedback projections versus horizontal lateral connections within a cortical layer, represents a natural next step that DynVision is designed to facilitate.

The importance of these often-implicit choices is further underscored by our reimplementation of CORnet-RT [15] and CordsNet [19] within DynVision. Both reimplementations revealed hidden design decisions such as initial idle periods, loss computation windows, and implicit feedforward delays that substantially affect learned dynamics but were not explicitly discussed in the original publications. For example, CORnet-RT’s classification capability collapses when even a single null-input timestep precedes the stimulus, a property that is a direct consequence of its recurrence delay configuration rather than a principled limitation of its architecture. These observations reinforce the need for frameworks that make all modeling choices visible and systematically explorable.

The current work has several limitations that influence interpretation. First, all demonstration results use ImageNette, a 10-class subset of ImageNet. While sufficient for characterizing temporal dynamics and comparing recurrence types, generalization of these findings to larger-scale classification tasks remains to be validated. Second, the framework implements rate-based dynamics, which capture population-level temporal phenomena but cannot model spike-timing-dependent effects or the computational effects of discrete action potentials. Third, training relies on standard backpropagation through time, which scales memory linearly with sequence length; alternative credit assignment methods that avoid storing the full computational graph could extend the feasible range of simulation durations [63], enabling investigation of dynamics over behaviorally relevant timescales of hundreds of milliseconds while remaining computationally efficient. Finally, our temporal step size of *dt* = 2 ms is arbitrary with respect to absolute time, and the correspondence between model timesteps and biological time should be treated as a modeling convention rather than a claim about precise temporal alignment with cortical dynamics.

Taken together, our results address several of the open questions that motivated this work. Biologically plausible temporal dynamics can emerge from simple recurrent connections combined with metabolic constraints, without requiring explicit normalization mechanisms. Different connectivity configurations give rise to functionally distinct dynamic regimes, demonstrating that not all recurrence is computationally equivalent. And these biological properties can be achieved without sacrificing computational efficiency. Looking forward, we intend to extend the framework to support excitatory-inhibitory cell-type separation, cell-type and hierarchy-specific connectivity rules [64, 65, 66], biologically plausible learning rules, unsupervised (contrastive) loss functions [67], attentional gain control [21], and elaborating our implementations of supralinear activation functions and pre-cortical processing (Supplementary Section 8.2) enabling more direct correspondence between model components and cortical circuit elements. By making the space of biologically plausible RCNN architectures systematically explorable, DynVision provides a foundation for resolving open questions about the functional roles of recurrence in visual processing, and for building models that are simultaneously performant, interpretable, and biologically grounded.

## 5 Code and Data Availability

The DynVision toolbox is available as open-source software at https://github.com/Lindsay-Lab/ DynVision under the MIT License, with comprehensive documentation at https://lindsay-lab.github.io/dynvision/. The current release (v0.1) is archived with a permanent DOI via Zenodo. Example configurations, pre-trained model weights (hosted on Hugging Face at https://huggingface.co/neurograce/DynVision), and the processed output data supporting all results presented in this paper are included in the repository.

All results in this work are based on models trained with at least three different random initialization seeds. The fully resolved YAML configuration files for every trained model are archived alongside the model checkpoints, ensuring complete reproducibility of all experimental conditions. The toolbox is actively maintained with semantic versioning and backward compatibility policies. Community contributions are welcomed through GitHub issues and pull requests.

The training data used in this study (ImageNette) is publicly available. For custom datasets, instructions for integrating new data sources are provided in the documentation.

## 6 Acknowledgments

We thank Wayne Soo for helpful discussions about training recurrent networks.

## 8 Supplementary Information

### 8.1 Software Architecture Details

This section provides detailed descriptions of the core software components summarized in the main Methods Section 2.6 (Software Design). Each subsection elaborates on the implementation choices and internal structure of one component of the toolbox.

#### 8.1.1 Base Classes

DynVision employs a modular base class architecture that separates concerns into specialized components, each addressing distinct aspects of neural network modeling. This design leverages multiple inheritance and mixin classes so that a `BaseModel` integrates all components, while researchers can also compose models from a subset of base classes or add custom parent classes with specific functionality.

##### TemporalBase

The `TemporalBase` class manages the time dimension of signal tensors (`[batch_size, n_timesteps, n_channels, dim_y, dim_x]`), including temporal unrolling and residual timestep handling. Its `forward()` method processes input sequentially across time; at each timestep, `_forward()` iterates over the layer names and executes each layer’s operations in the order defined by its `layer_operations` list (see Section 2.3.4). Any operation not defined for a given layer is silently skipped, enabling heterogeneous layer configurations within a single model.

##### LightningBase

The `LightningBase` class integrates PyTorch Lightning [48] to handle the engineering overhead of model training: automatic mixed-precision training, structured logging, checkpointing, and hardware acceleration from single GPUs to multi-node clusters. It provides a Lightning module preconfigured for training recurrent convolutional networks, with hyperparameters such as learning rate, optimizer choice, and scheduler type set to reasonable defaults and overridable via the configuration files.

##### StorageBuffer

The `StorageBuffer` component addresses the memory demands of recording model outputs and layer activations selectively during training, validation, or testing and across potentially hundreds of timesteps. It supports fixed and cyclic recording strategies with configurable buffer sizes and optional CPU offloading to capture population dynamics without exhausting GPU memory. The recorded data can be exported as tensors, dictionaries, or pandas DataFrames combining labels, predictions, batch and sample indices, timesteps, and per-unit activations to enabling direct comparison with electrophysiological recordings that preserve the full temporal evolution of neural activity.

##### DtypeDeviceCoordinator

While PyTorch Lightning handles most device and dtype coordination, the `DtypeDeviceCoordinator` provides an additional safety layer: it ensures consistency across child and parent classes (which may not all derive from `LightningModule`), variables in storage buffers, and between the dataloader and model trainer.

##### Monitoring

The `Monitoring` class provides structured logging of model, training, and system parameters, including detailed GPU and CPU memory usage. It automatically detects common numerical issues such as NaN gradients and parameter drift, tracks parameter evolution across epochs, and supports real-time visualization of network dynamics and weight distributions. The logging integrates with PyTorch Lightning’s callback system and external tools such as Weights&Biases ^2^.

#### 8.1.2 Parameter Handling

##### Hierarchical Configuration

The toolbox implements a comprehensive parameter management framework that separates configuration from code, enhancing reproducibility and systematic exploration of biological parameter spaces. The system employs human-readable YAML configuration files organized hierarchically across specialized domains: `config_defaults.yaml` establishes baseline model parameters and system settings, `config_data.yaml` defines dataset specifications, `config_experiments.yaml` specifies data selection and parameter sweep protocols for model testing scenarios. The `config_workflow.yaml` file sits at the top of the hierarchy and defines parameters relevant for the current application, overwriting duplicate default settings in other config files. Further, `config_modes.yaml` provides parameter presets to rapidly switch between configuration contexts such as `local` for local test runs, `debug` for verbose logging, `large_dataset` for memory-optimized processing, and `distributed` for multi-device cluster execution. This hierarchical organization enables researchers to override specific parameters at appropriate abstraction levels while maintaining consistent defaults.

##### Parameter Precedence

The system implements a three-level precedence hierarchy that provides precise control over parameter resolution: source precedence (command-line arguments override configuration files), scope precedence (component-specific parameters like `model.learning_rate` override global defaults), and alias precedence (short-form aliases override their long-form equivalents at the same scope level). This precedence system ensures that researchers can override specific parameters at appropriate abstraction levels while maintaining parameter transparency and predictability across different specification methods.

##### Adaptive Mode System

The framework includes an auto-detection mode system that adapts parameters to execution contexts without manual configuration. The `ModeRegistry` automatically detects scenarios such as debug mode (verbose logging for development), large dataset mode (memory-optimized settings for high-resolution or long-sequence data), and distributed mode (multi-device coordination), adjusting parameters accordingly while allowing explicit user overrides. This mode system reduces configuration burden while maintaining reproducibility, as active modes are recorded in the provenance metadata.

##### Parameter Expansions and Overwrites

The Snakemake workflow system extends this configuration framework through dynamic parameter expansion and command-line integration. Configuration parameters can be selectively overridden via command-line arguments (`snakemake –config model_name=DyRCNNx4 lr=0.001`), enabling rapid prototyping without modifying configuration files. Parameter lists are automatically expanded into Cartesian products for systematic exploration (`recurrence_type: [full, self, depthwise]` generates separate jobs for each architecture variant), while runtime configurations are automatically archived alongside experimental outputs to ensure complete reproducibility.

##### Parameter Parsing

The script-level parameter handling employs type-safe Pydantic classes that provide robust validation, automatic type conversion, and biological constraint checking. Specialized parameter classes (`ModelParams`, `DataParams`, `TrainerParams`) encapsulate domain-specific validation logic and computed properties, while composite classes (`InitParams`, `TrainingParams`, `TestingParams`) combine these components for script-specific processing. The system supports flexible parameter aliasing (`lr` → `learning_rate`, `rctype` → `recurrence_type`) and implements context-aware validation that enforces system and biological feasibility constraints and cross-parameter consistency checks, e.g., time delays are integer multiples of time resolution, or batch size and learning rate scaling for multi-device training and gradient accumulation.

##### Reproducibility Through Configuration Persistence

To ensure complete reproducibility, the parameter system persists the fully resolved configuration alongside each experimental output. Every model check-point, response export, and analysis result is accompanied by a `<artifact>.config.yaml` file containing the complete parameter specification, active modes, and provenance metadata indicating whether each parameter value originated from configuration files, mode payloads, or command-line overrides. This practice ensures that every artifact carries its own reproducibility record, enabling precise replication of experimental conditions and transparent documentation of the parameter choices underlying each result.

#### 8.1.3 Data Handling

The data management pipeline (summarized in the main Methods **??**) is implemented as a Snakemake-managed workflow that transforms raw datasets into model-ready formats.

##### Dataset Partitioning and Subsetting

Raw datasets are organized into training and testing splits via a symbolic-link-based directory structure rather than file duplication, eliminating storage overhead when defining overlapping subsets for different experiments. Class subsets are specified as index lists in `config_data.yaml` (e.g., `class_subset: [0, 5, 12, 28]`). The pipeline automatically resolves label-index transformations: when a model trained on the full dataset is evaluated on a subset, the system remaps class indices to maintain classifier-output alignment, and when a model is trained on a subset, its classifier layer is automatically resized to match the number of presented classes.

##### Handling Non-Standard Dataset Layouts

For datasets where on-disk folder names do not correspond to sequential numeric class labels (e.g. ImageNet’s identifiers like `n01440764`) the toolbox accepts a JSON mapping file that translates between directory names and zero-indexed class labels. This mapping is applied once during symbolic-link construction; all downstream operations (data loading, training, evaluation) then use uniform numeric indexing, making it straightforward to train on one dataset and test on a class subset of another.

##### FFCV Integration

For temporal models that process image sequences over many timesteps, the CPU-to-GPU transfer of per-timestep image batches can dominate training time. The toolbox integrates FFCV (Fast Forward Computer Vision) [27] as an optional dataloader backend. FFCV pre-processes and stores datasets in a compressed, memory-mapped binary format that enables direct GPU-copy without Python-level deserialization. Temporal expansion (duplicating a static image across timesteps) can be performed either during dataset construction (FFCV pipeline, minimizing per-epoch cost) or at runtime (model forward pass, enabling dynamic pattern variation). Benchmark comparisons between FFCV and standard PyTorch DataLoaders are provided in Section 3.9.

#### 8.1.4 Workflow Management

The workflow system (summarized in the main Methods) is built on Snakemake [49] and organized as a directed acyclic graph of rules.

##### Rule Structure

Each processing stage (e.g., data acquisition, dataset organization, model training, analysis, and figure generation) is defined as a Snakemake rule with explicit input and output file declarations. Rules are triggered only when their outputs are missing or when upstream inputs have changed, avoiding redundant recomputation. Parameter sweeps are expressed as YAML lists in configuration files (e.g., `recurrence_type: [full, self, depthpointwise]`), which Snakemake expands into independent jobs via its wildcard mechanism. Each job receives a fully resolved parameter snapshot written alongside its outputs, ensuring traceability from configuration to result.

##### Resource Management and Scaling

Resource requests (CPU cores, GPU devices, memory, runtime) are specified per rule and can be overridden via profiles for different execution environments. On a single machine, Snakemake parallelizes independent jobs up to the available core count. On HPC clusters, the toolbox provides SLURM submission profiles that translate rule resource declarations into scheduler directives, enabling automatic distribution of parameter sweeps across nodes. A centralized `config_paths.yaml` file resolves output directories, data locations, and checkpoint paths relative to the execution environment, so the same workflow definition runs identically on a laptop and a cluster without path modifications.

### 8.2 Additional Biological Components

#### Supralinearity

Individual neurons often exhibit nonlinear amplification, where strong inputs lead to disproportionately large activations. The influence of recurrence is necessary to stabilize this explosive activity [68]. The toolbox allows users to apply a supralinear activation function inspired by models of visual cortex [61, 69] to encourage networks to make effective use of recurrent connections for regularizing feedforward activity.

#### Pre-cortical Processing

Having a dedicated input layer to preprocess the input stimuli is a common component of vision models. The toolbox includes two biologically inspired versions of this: 1.) a simple adaptation factor (fixed or learnable) that reduces the input values to the network over time so that it is necessarily more dependent on recurrent connections to retain information, 2.) a layer representing the retina and LGN and their representational bottleneck, as described in [70].

### 8.3 Comparison to Neural Data

**Figure 15:**
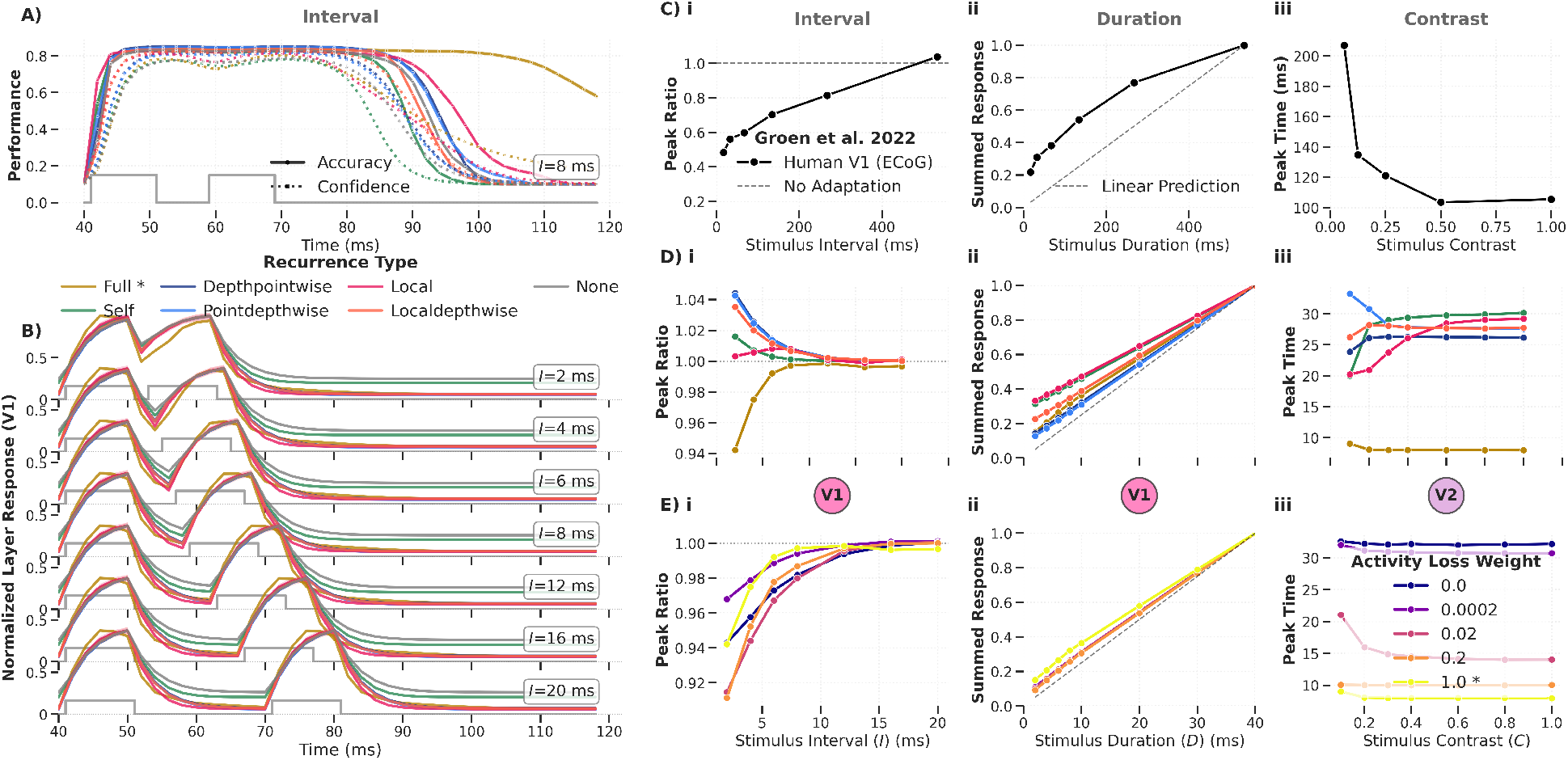
Temporal RCNN dynamics for different recurrence types in response to input with varied interval, duration, and contrast. Same as Figure 12 with layer summaries switched between V1 and V2.

**Figure 16:**
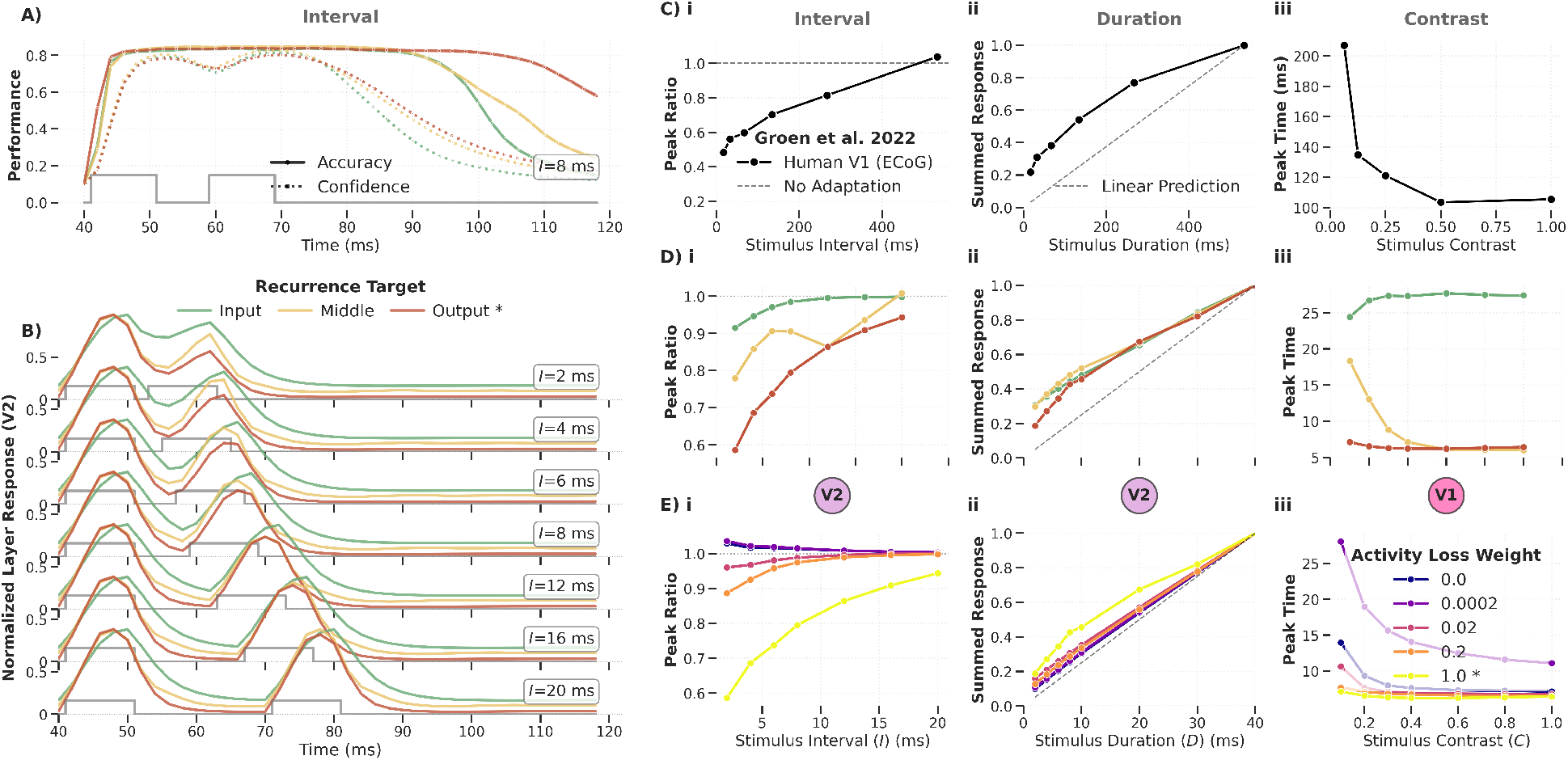
Temporal RCNN dynamics for different recurrence targets in response to input with varied interval, duration, and contrast. Same as Figure 12 but with recurrence type `full` but different recurrence targets.

**Figure 17:**
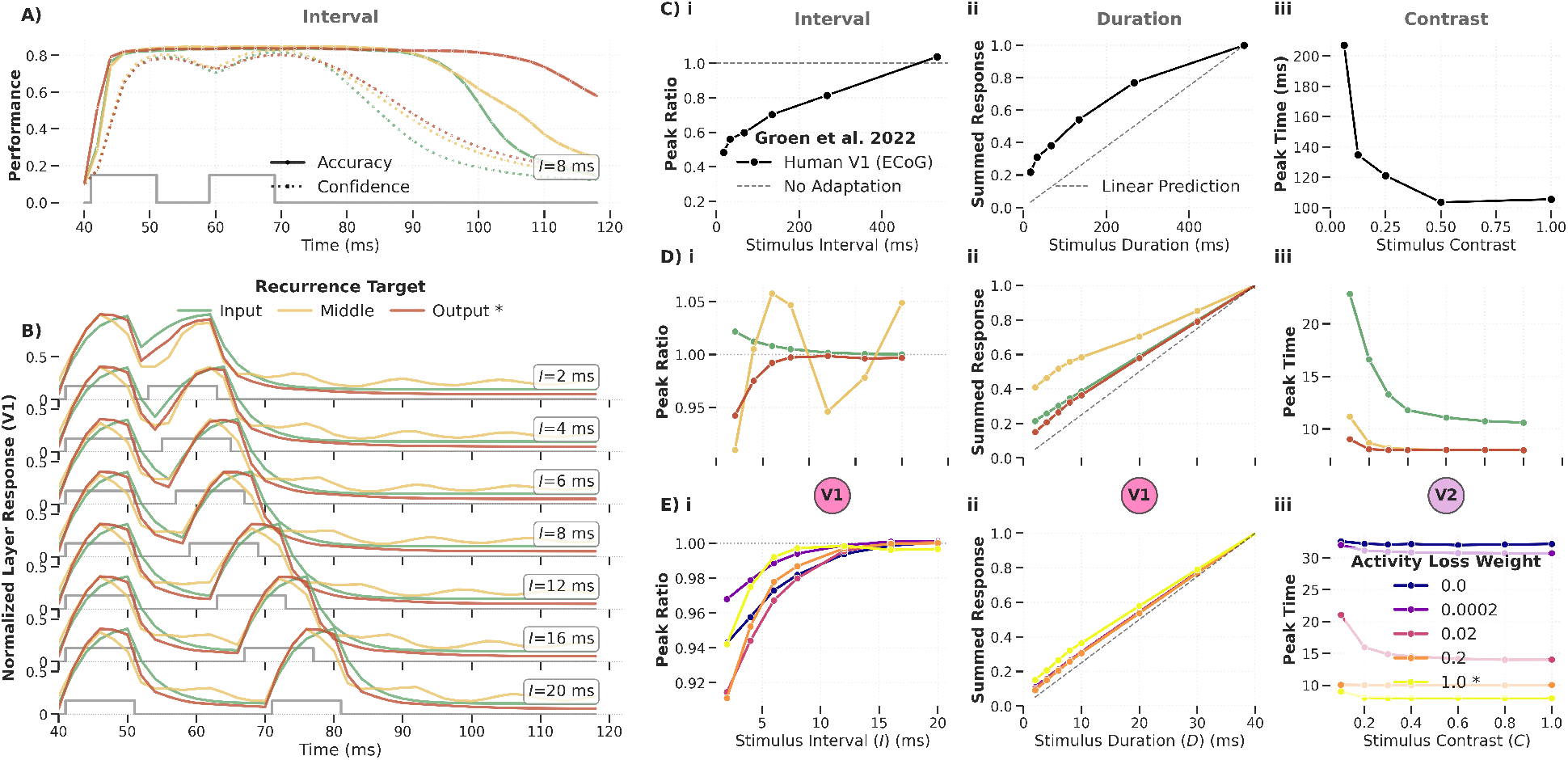
Temporal RCNN dynamics for different recurrence targets in response to input with varied interval, duration, and contrast. Same as Supplementary Figure 16 with layer summaries switched between V1 and V2.

### 8.4 Robustness to Noise

**Figure 18:**
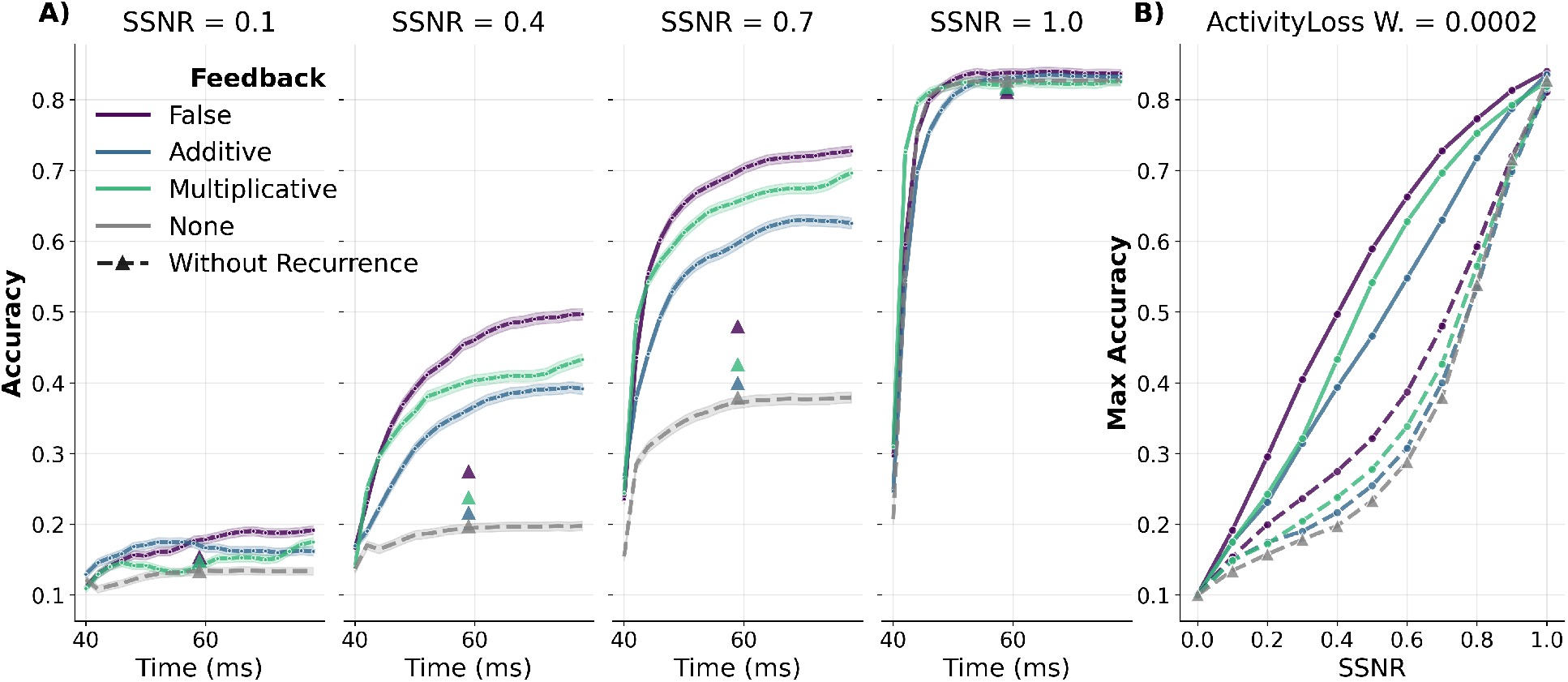
Noise robustness with feedback connections. Models with additive and multiplicative feedback connections (middle-target, full recurrence, activity loss weight = 0.0002) are tested on images with increasing Gaussian noise (SSNR). Across seeds, neither feedback type provides a consistent accuracy advantage over the no-feedback baseline, with multiplicative feedback being only marginally preferable to additive in some seeds.

**Figure 19:**
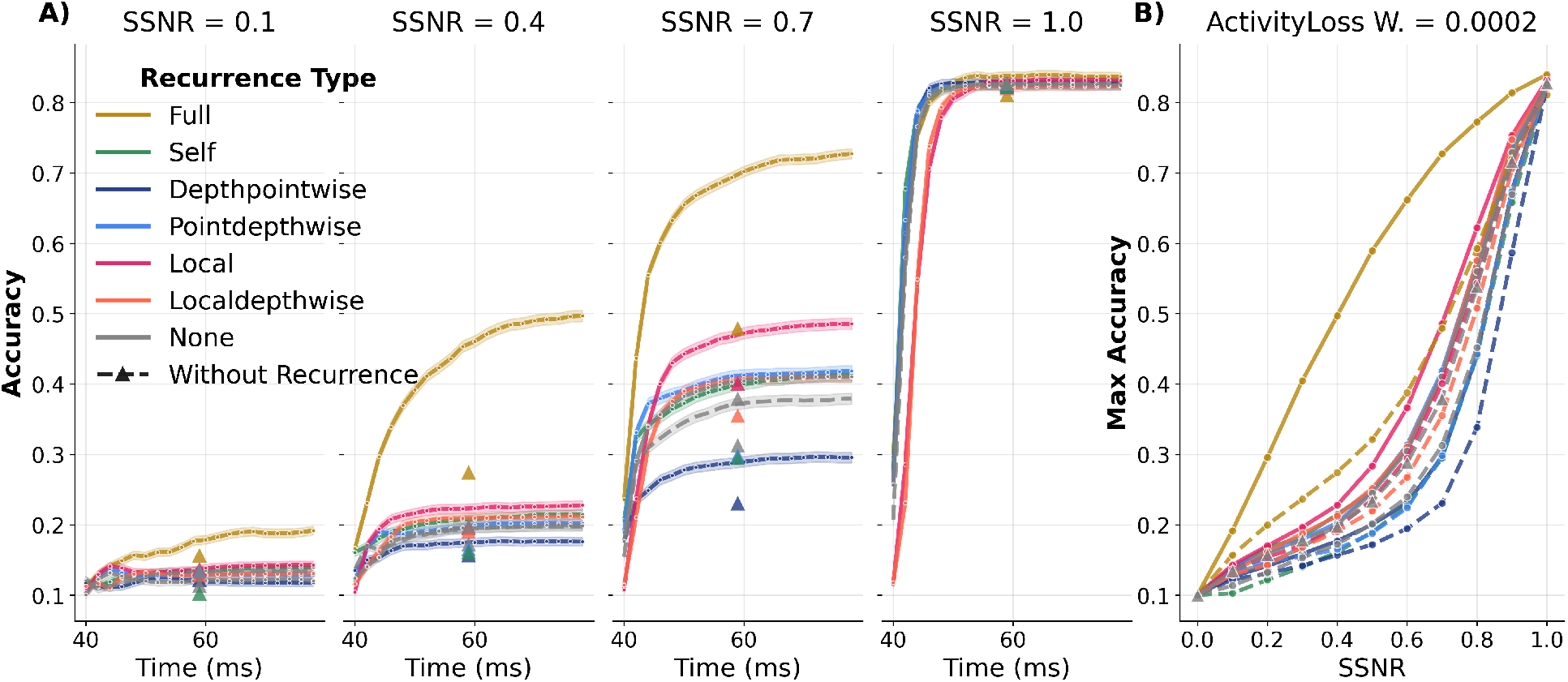
Noise robustness for different types of recurrence. Maximum accuracy as a function of noise intensity (SSNR) for models with recurrence target = middle, activity loss weight = 0.002, tested with different lateral recurrence types. Under these conditions, full recurrence outperforms depthpointwise, pointdepthwise, self, and local variants, consistent with the main-text finding that noise robustness depends jointly on recurrence type, integration target, and activity loss weighting.

### 8.5 Weight Distributions

**Figure 20:**
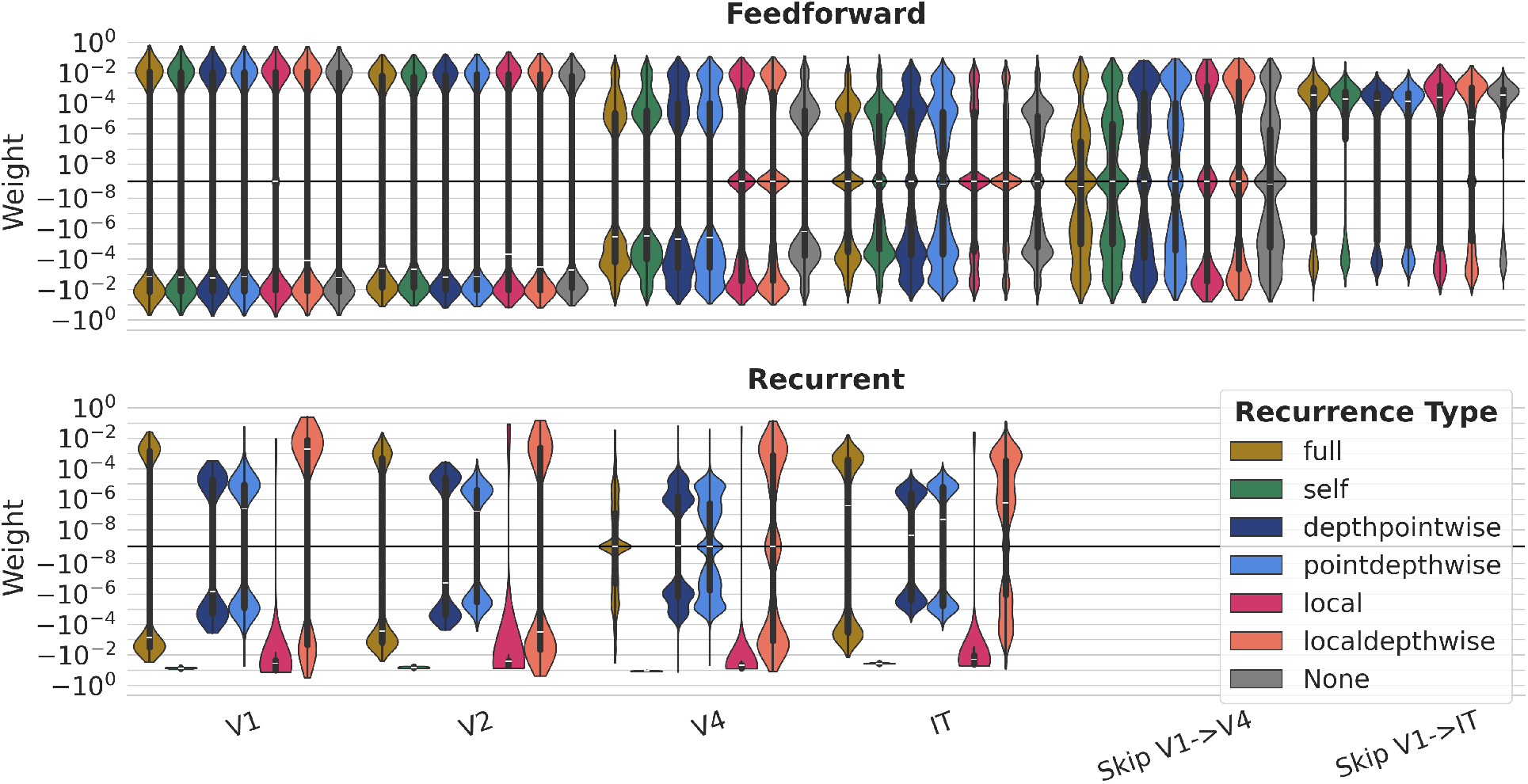
Weight distributions for models trained with different recurrence connectivity types. Feed-forward weights are consistent across all recurrence types (V1: *σ* ≈ 0.037; V2 *σ* ≈ 0.017; V4: *σ* ≈ 0.008; IT *σ* ≈ 0.004). Recurrent weight means vary markedly with connectivity type: self-recurrence produces single strongly negative weights around (−0.073 ± 0.008), local recurrence produces a distribution of mostly negative weights (−0.039 ± 0.031), full recurrence is weakly negative (−0.0004 ± 0.0024), and sparser types (depthpointwise, pointdepthwise) remain near zero with 0.000 ± 0.001 and 0.000 ± 0.002 respectively, localdepthwise recurrence is the only leaning slightly positive (0.0002 ± 0.0095).

**Figure 21:**
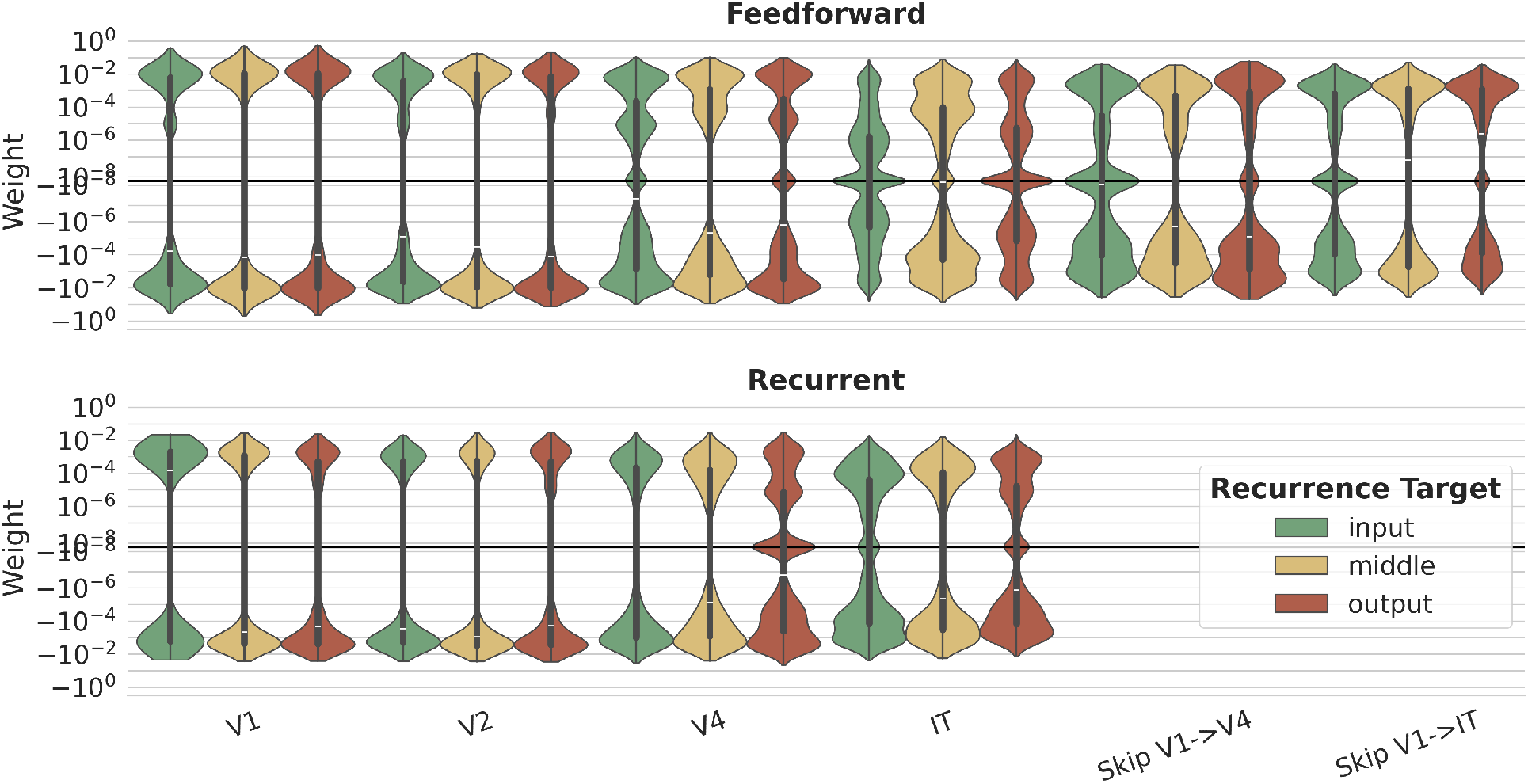
Weight distributions for models trained with different recurrent connection targets. Feed-forward weight distributions are consistent across recurrence targets (V1: *σ* ≈ 0.031; V2: *σ* ≈ 0.018; V4: *σ* ≈ 0.009; IT: *σ* ≈ 0.004). Recurrent weights are mildly negative across all targets with comparable magnitudes: input (−0.0003 ± 0.0027), middle (−0.0006 ± 0.0028), output (−0.0006 ± 0.0029).

**Figure 22:**
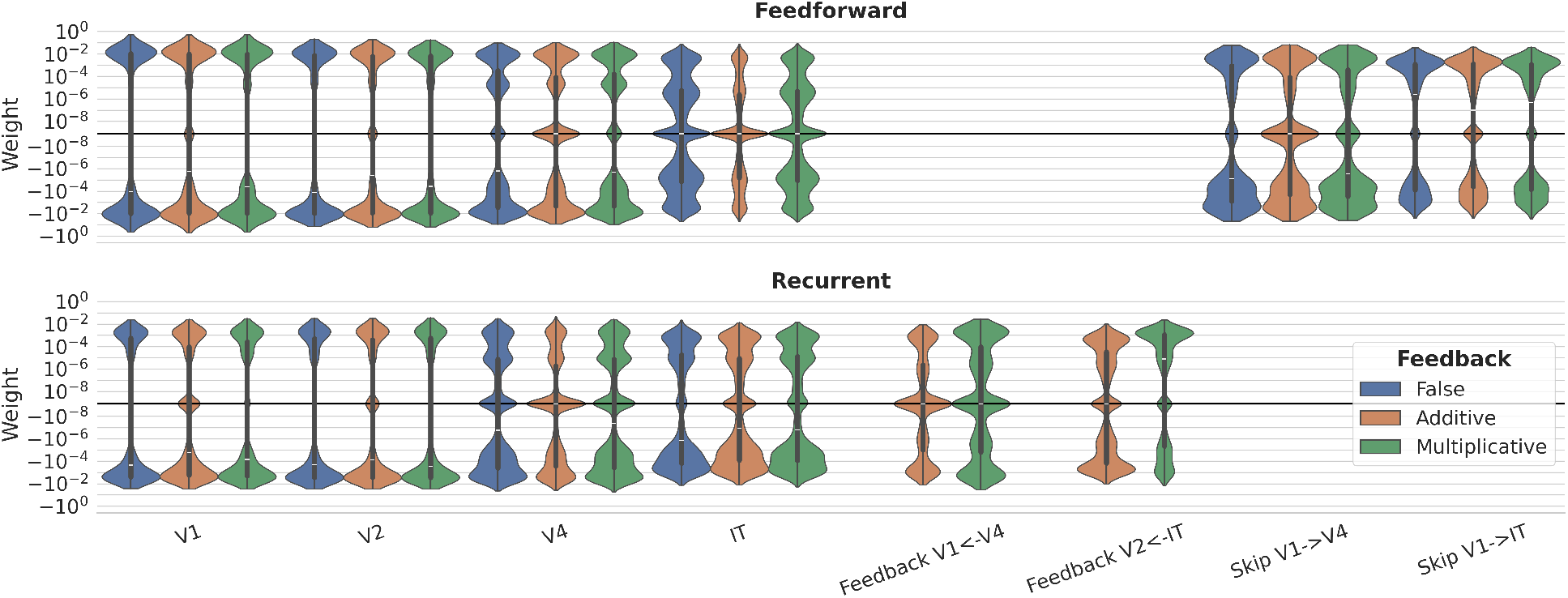
Weight distributions for models trained with different feedback connection types. Feedforward weights (V1: 0.0007 ± 0.035; V2: 0.0 ± 0.0196; V4: −0.0005 ± 0.010; IT: 0.0 ± 0.004) and recurrent weights (V1: −0.001 ± 0.003; V2: −0.001 ± 0.004; V4: −0.001 ± 0.003; IT: 0.0 ± 0.001) are unaffected by the feedback condition. Feedback weights remain near-zero after training: additive feedback (−0.00005 ± 0.0009) is essentially unchanged from its near-zero initialization (0.0 ± 0.001), while multiplicative feedback shows only a modest positive shift (+0.0004 ± 0.0023).

**Figure 23:**
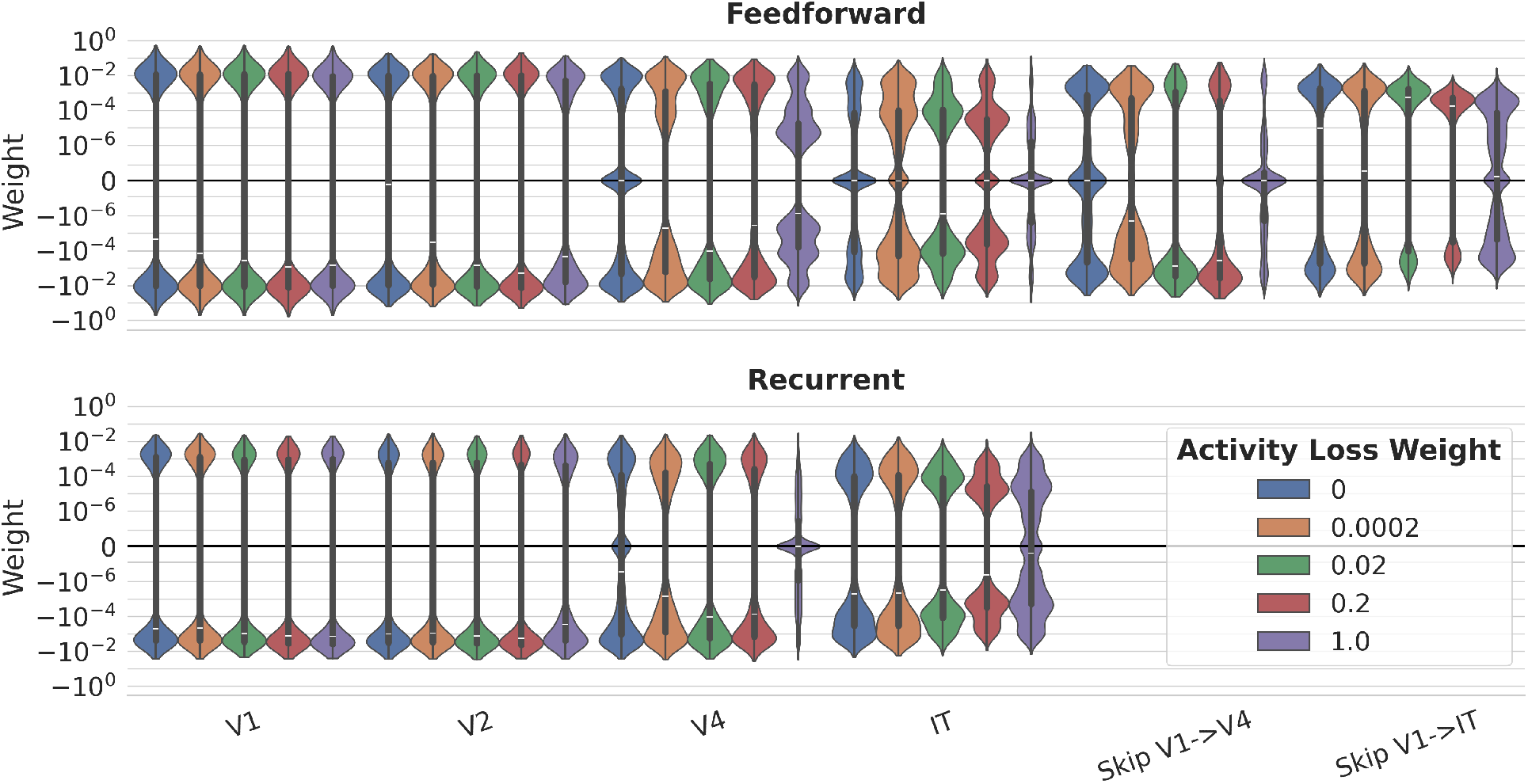
Weight distributions for models trained with different activity loss factors (pattern = 1). Feedforward weight distributions are stable across all tested regularization strengths (V1: std ≈ 0.034; V4: std ≈ 0.010). Recurrent weights remain consistently small and negative across all activity loss values (*λ*_act_ = 0-1.0), with per-condition means ranging from −0.00064 to −0.00098 and std ≈ 0.0025-0.0030.

**Figure 24:**
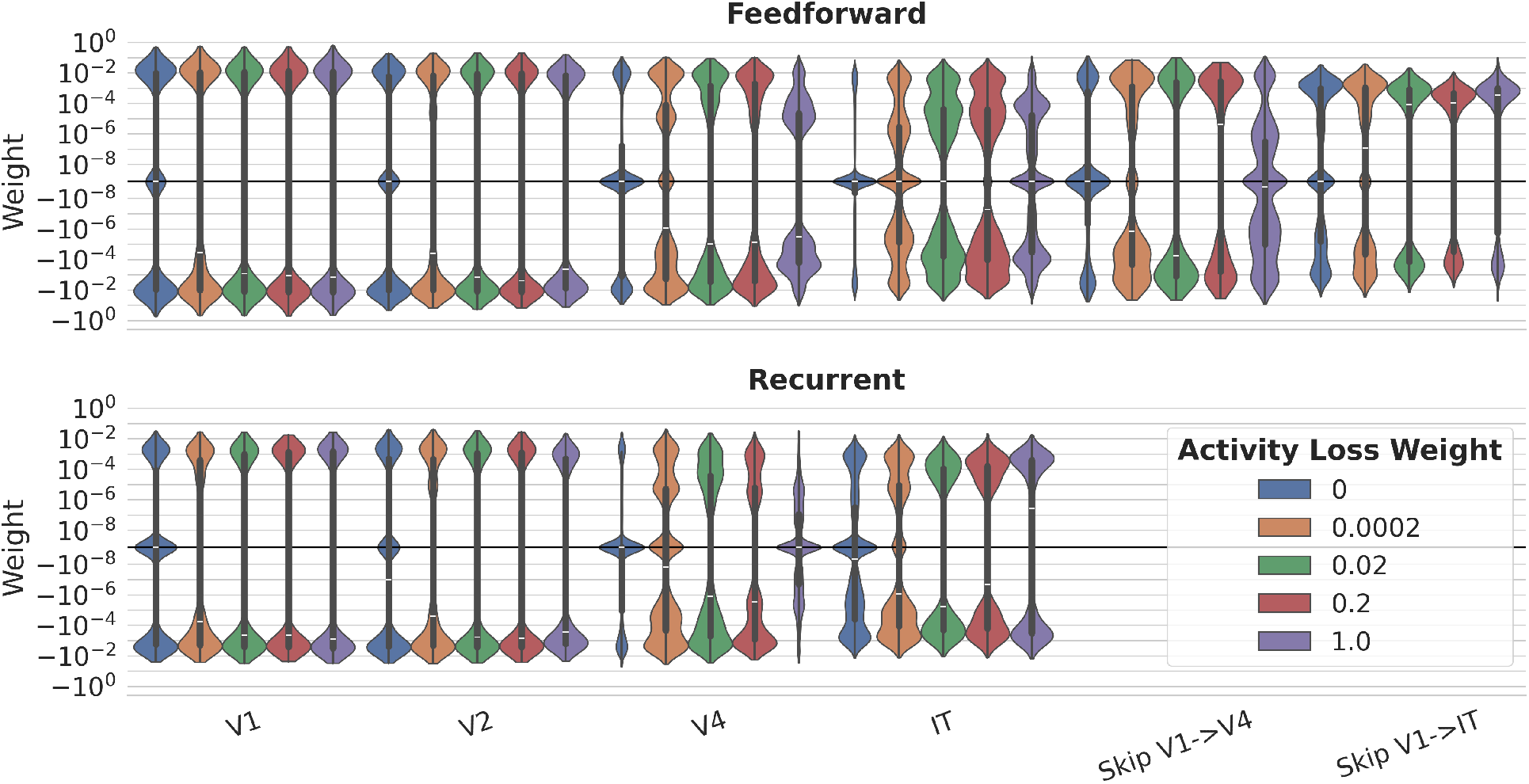
Weight distributions for models trained with different activity loss factors (pattern = 1011). Results are qualitatively consistent with pattern = `1` (Supplementary Figure 23). Feedforward weights are unaffected by regularization strength (V1: std ≈ 0.037; IT: std ≈ 0.004). Recurrent weights remain small and negative across all activity loss values, with per-condition means ranging from −0.00039 to −0.00050 and std ≈ 0.0024-0.0028.

### 8.6 Stability Evaluation

**Figure 25:**
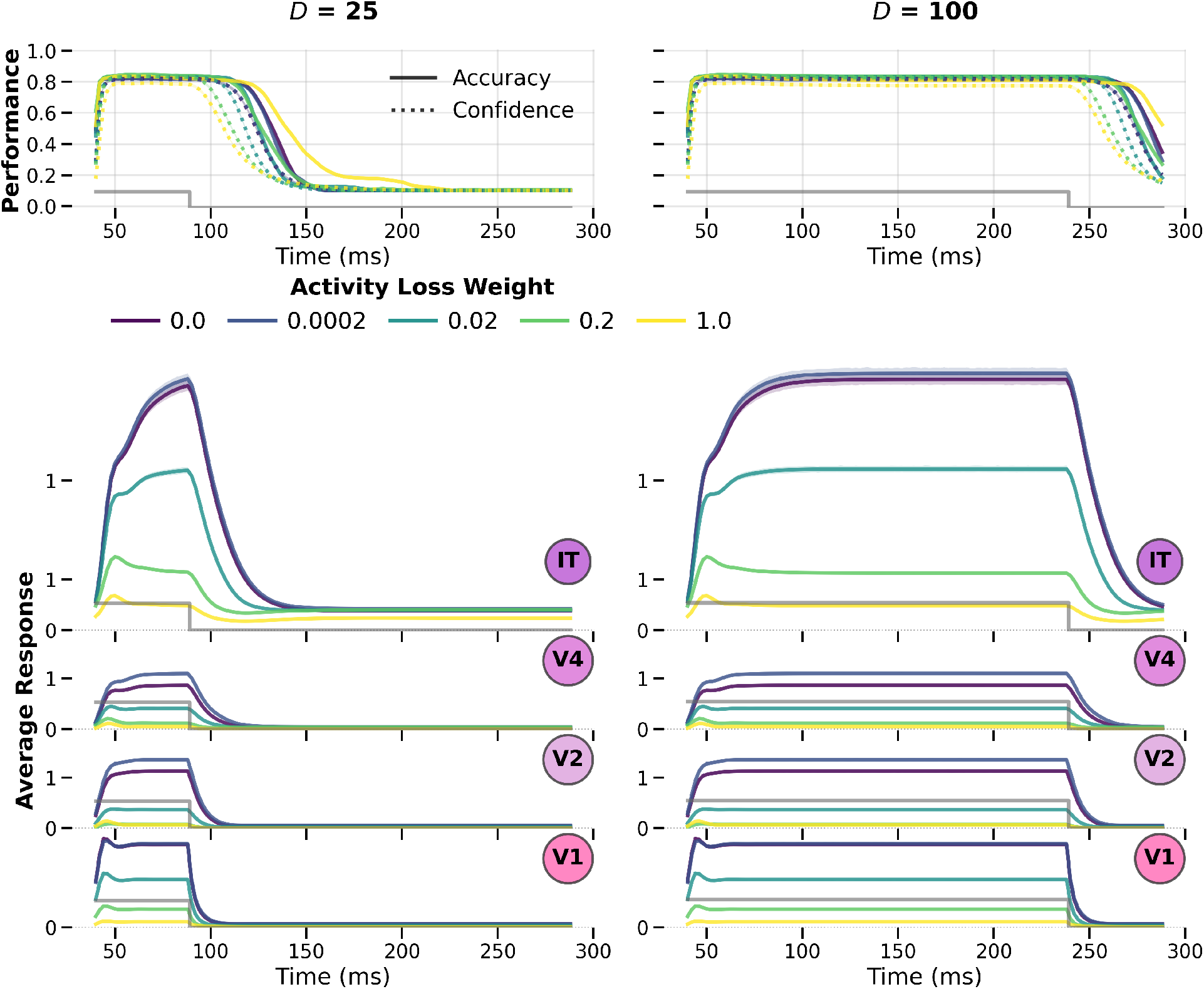
Models exhibit stable dynamics and appropriate null responses. Same as Figure 11 but for model variations trained with interspersed image presentation pattern = `1011`.

**Figure 26:**
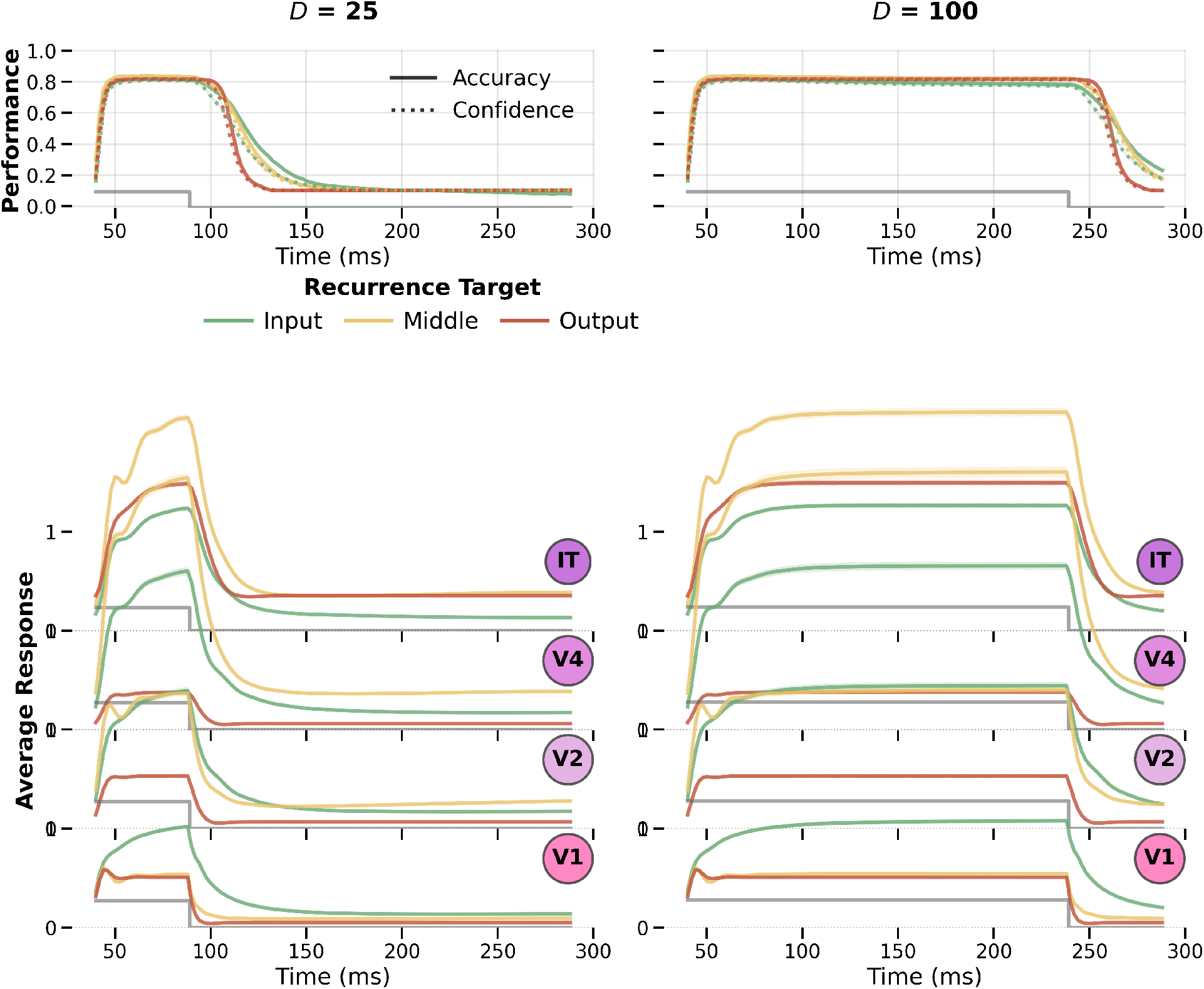
Models exhibit stable dynamics and appropriate null responses. Same as Figure 11 but for model variations trained with activity loss weight = 0.0002 and different recurrence targets.

### Footnotes

1 https://docs.pydantic.dev/latest/

2 https://wandb.ai

